# Resolving the three-dimensional interactome of Human Accelerated Regions during human and chimpanzee neurodevelopment

**DOI:** 10.1101/2024.06.25.600691

**Authors:** Atreyo Pal, Mark A. Noble, Matheo Morales, Richik Pal, Marybeth Baumgartner, Je Won Yang, Kristina M. Yim, Severin Uebbing, James P. Noonan

**Author notes:** Correspondence to: James P. Noonan.

## Abstract

Human Accelerated Regions (HARs) are highly conserved across species but exhibit a significant excess of human-specific sequence changes, suggesting they may have gained novel functions in human evolution. HARs include transcriptional enhancers with human-specific activity and have been implicated in the evolution of the human brain. However, our understanding of how HARs contributed to uniquely human features of the brain is hindered by a lack of insight into the genes and pathways that HARs regulate. It is unclear whether HARs acted by altering the expression of gene targets conserved between HARs and their chimpanzee orthologs or by gaining new gene targets in human, a mechanism termed enhancer hijacking. We generated a high-resolution map of chromatin interactions for 1,590 HARs and their orthologs in human and chimpanzee neural stem cells (NSCs) to comprehensively identify gene targets in both species. HARs and their chimpanzee orthologs targeted a conserved set of 2,963 genes enriched for neurodevelopmental processes including neurogenesis and synaptic transmission. Changes in HAR enhancer activity were correlated with changes in conserved gene target expression. Conserved targets were enriched among genes differentially expressed between human and chimpanzee NSCs or between human and non-human primate developing and adult brain. Species-specific HAR gene targets did not converge on known biological functions and were not significantly enriched among differentially expressed genes, suggesting that HARs did not alter gene expression via enhancer hijacking. HAR gene targets, including differentially expressed targets, also showed cell type-specific expression patterns in the developing human brain, including outer radial glia, which are hypothesized to contribute to human cortical expansion. Our findings support that HARs influenced human brain evolution by altering the expression of conserved gene targets and provide the means to functionally link HARs with novel human brain features.

## Introduction

The human brain exhibits distinct features compared to the brains of other primate species, including increased cortical surface area, a larger number of neurons, and prolongation of neurogenesis and synaptogenesis^1,2^. Genetic changes that altered the level, timing and spatial distribution of gene expression during human evolution are hypothesized to contribute to these developmental differences^3,4^. One prominent class of elements, termed Human Accelerated Regions (HARs), have been implicated in the evolution of novel human traits, particularly in the brain. HARs are highly conserved across species but show accelerated evolution on the human lineage, suggesting they may have gained novel functions^5–9^. Multiple studies support that HARs include transcriptional enhancers with human-specific activity^10,11^. HARs have been shown to drive changes in gene expression in genetically modified mouse models^12–15^, while massively parallel reporter assays (MPRAs) comparing HARs with their non-human primate orthologs have identified HARs that exhibit species-specific changes in enhancer activity^16–18^. Targeted perturbations and large-scale CRISPR proliferation screens have also been used to link HARs to changes in gene expression and neural stem cell phenotypes^19,20^. Another class of regulatory elements with a potential role in human brain development and evolution, known as Human Gained Enhancers (HGEs), have been identified based on increased levels of histone modifications associated with enhancer activity in the developing human brain compared to other species^21–23^. A subset of HGEs show altered activity in mouse models or in MPRAs^16,24^. However, insight into the role of most HARs and HGEs in human neurodevelopment remains limited.

HARs and HGEs may drive changes in gene expression via two non-exclusive mechanisms. First, HARs, HGEs and their chimpanzee orthologs may target conserved sets of orthologous genes in each species, with HARs and HGEs acting to alter expression of those genes in human^12–15,24^. Alternatively, it has been suggested HARs may have gained new gene targets in human neurodevelopment, a phenomenon referred to as enhancer hijacking^7^. Understanding the role of HARs and HGEs in human evolution as well as their predominant mechanism of regulation requires empirical insight into the gene target repertoire of HARs, HGEs and their chimpanzee orthologs and the pathways and cell types in which those genes operate. Machine learning approaches have been used to predict that 31% of HARs are putative enhancers and to infer their targets based on epigenetic data from multiple cell and tissue types^18^. However, these *in silico* approaches have yielded sets of gene target predictions that must then be experimentally validated. Recent studies have also identified gene targets for some HARs and HGEs by using chromosome conformation capture methods such as HiC in regions of the developing human brain^21,25,26^, as well as in cultured neural progenitors^7,27^. These targets include genes involved in neurodevelopmental processes such as neurogenesis and axon guidance, and genes implicated in autism spectrum disorder (ASD) and schizophrenia^26,28^. Collectively, these studies have identified potential targets for only 6%-33% of HARs due to the sparsity of HiC datasets (Table 1), which makes precise identification of gene targets challenging^7,18,26,28^. The majority of HARs and HGEs therefore have yet to be experimentally linked to specific genes, and potential interactions between HARs and HGEs and other regulatory elements remain uncharacterized. This also limits insight into whether HARs and HGEs share gene targets in human and chimpanzee or frequently engage in enhancer hijacking events.

**Table 1:**
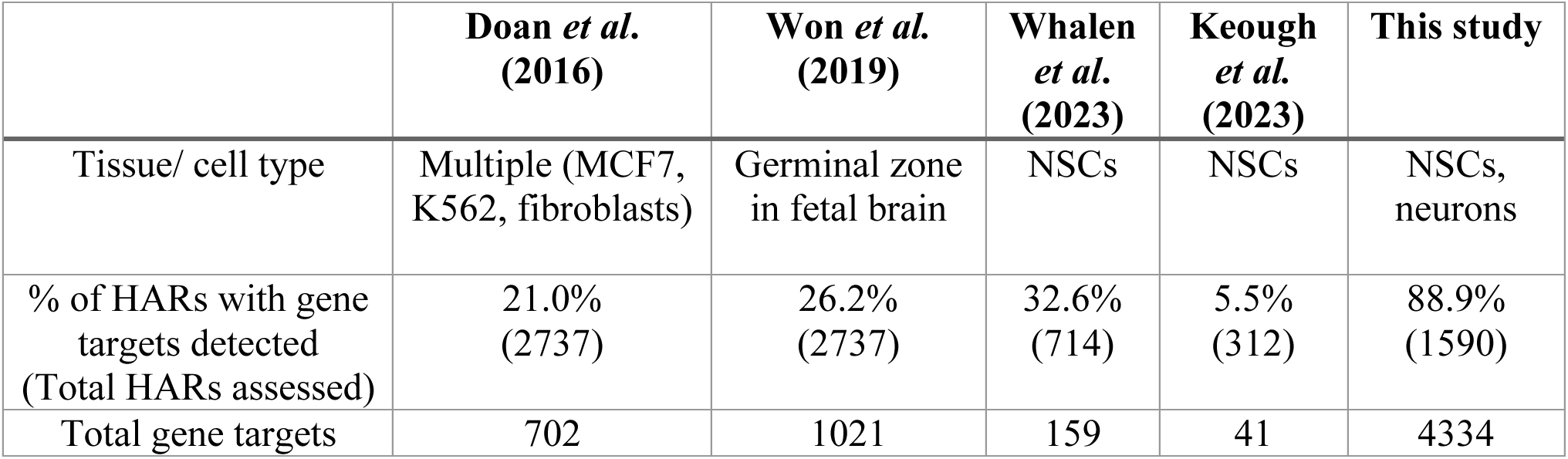
Comparison of this study with prior HiC studies linking HARs to genes. The total number of HARs considered in each study is shown by the numbers in parentheses below the proportion of HARs with assigned gene targets. See Methods for details.

To overcome these obstacles, we employed Capture HiC (CHi-C), which allows chromatin interactions involving thousands of genomic loci to be characterized at high resolution in a single experiment^29–31^. We applied this approach in human and chimpanzee iPSC-derived neural stem cells (NSCs) to define interactions for 1,590 HARs, as well as for 466 HGEs that exhibit differential enhancer activity compared to their chimpanzee orthologs^5,6,8,9,16^. We identified gene targets for 88.9% of HARs and 75.8% of HGEs in human NSCs. Comparing gene targets in human and chimpanzee identified a set of conserved genes targeted by HARs and HGEs and their chimpanzee orthologs, as well as species-specific gene targets. Conserved gene targets converged on biological processes related to neuronal migration, axon guidance and synaptic transmission, while species-specific targets did not converge on any known function. To complement these dense interaction maps, we generated maps of gene expression, histone modifications associated with active or repressed chromatin states, chromatin looping factor binding events, and RNA Polymerase II binding to reveal interactions among HARs and HGEs and other regulatory elements. These analyses also identified a subset of HAR-gene target interactions that switched from a repressed state in NSCs to an active state in neurons. Species differences in HAR and HGE activity were associated with differences in conserved gene target expression, and HAR and HGE conserved gene targets were significantly overrepresented among genes showing human-specific expression changes in the brain^32–34^. In contrast, species-specific targets were not overrepresented among differentially expressed genes. Integrating HAR and HGE target sets with a public human embryonic and fetal brain single-cell gene expression atlas^35,36^ identified genes showing biased expression in specific cell types. Together, our results support that HARs and HGEs are directly associated with changes in gene regulation via genes commonly targeted in both species during neurodevelopment and provide a comprehensive view of the biological processes and cell types that HARs and HGEs may have altered in human brain evolution.

## Results

### Mapping HAR and HGE interactions in human and chimpanzee neural stem cells

To generate maps of HAR and HGE chromatin interactions, we performed CHi-C in induced pluripotent stem cell (iPSC)-derived human and chimpanzee neural stem cells (hNSCs and cNSCs; Methods). Both the hNSCs and cNSCs used in our experiments expressed known NSC markers such as *NES, SOX2, PAX6,* and *FABP7* (Fig. S1A, S1C) and comprised a population of transcriptionally similar cells based on single-cell RNA sequencing (scRNA-seq) (Fig. S1B; Methods)^36–40^. Additionally, we demonstrated that these hNSCs could be differentiated into excitatory neurons characterized by the expression of markers including *TUJ1*, *MAP2, DCX, VGLUT1* and *VGLUT2* (Fig. S1C; Methods)^40–43^. We generated CHi-C maps for hNSCs and cNSCs by integrating three CHi-C replicates generated from independent NSC derivations in each species. We also generated and integrated two CHi-C replicates from independent differentiations to obtain interaction profiles in human neurons (Methods). In parallel, we generated gene expression profiles and maps of the histone modifications H3K27ac, which is associated with active promoters and enhancers, and H3K27me3, which is associated with repressed regions; binding events for CTCF and the cohesin complex subunit RAD21, which mediate regulatory interactions; and RNA Polymerase II in hNSCs, cNSCs and human neurons^44–46^. We integrated these functional genomic datasets with the interaction maps to provide insight into interaction regulatory states. A summary of the datasets generated in this study is provided in Table S1.

We first sought to characterize and compare interaction profiles for HARs, HGEs and their chimpanzee orthologs in NSCs. We identified a total of 39,093 interactions in hNSCs and 26,668 interactions in cNSCs, yielding interactions for > 90% of the HARs, HGEs and their orthologs that we targeted (Methods; Fig. 2A)^47–49^. Despite the overall difference in the number of interactions detected between human and chimpanzee, the proportions of conserved and species-specific interactions assigned to HARs and HGEs were similar (Fig. 2A; Methods). We found that 88.4% of HARs and 86.9% of HGEs have at least one interaction in common with their chimpanzee ortholog. The median proportions of conserved interactions for HARs and their chimpanzee orthologs were 62.1% and 70.9%, respectively (Fig. 2A). One example illustrating the similarity of interaction profiles for one HAR, *HACNS240*, and its chimpanzee ortholog is presented in Fig. 2B and Fig. S2A. This HAR was previously shown to exhibit higher enhancer activity than its chimpanzee ortholog in an MPRA study in human neural stem cells^16^. Our approach captures a high local density of significant interactions within a 600kb genomic interval around *HACNS240*. We then defined gene targets based on whether an interaction fell within an annotated gene or 5kb upstream of the annotated transcription start site (GENCODEv43 annotation; Methods). Based on these criteria, the identified gene targets of *HACNS240* and its chimpanzee ortholog, namely *DUSP6*, *POC1B*, *GALNT4* and *LINC02458*, are conserved. An additional example, HGE531, shows a similar pattern of gene target conservation (Fig. S2B). The extent of gene target conservation we observe in these examples suggests that HARs, HGEs and their orthologs may generally target orthologous genes.

**Figure 1:**
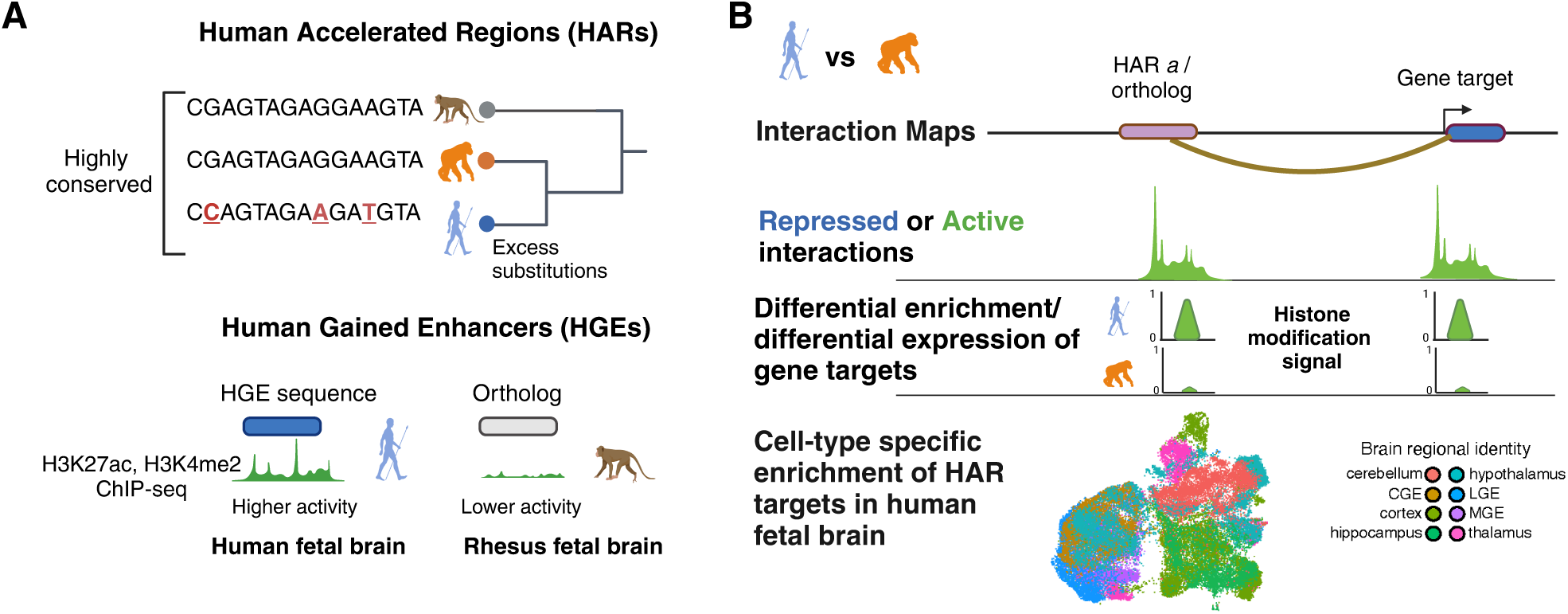
Overview of the study. **(A)** Schematic illustrating how HARs and HGEs are defined. **(B)** Outline of the datasets and analyses used to identify and annotate HAR and HGE interactions as described in the Results. Schematics were generated using BioRender.com.

**Figure 2.**
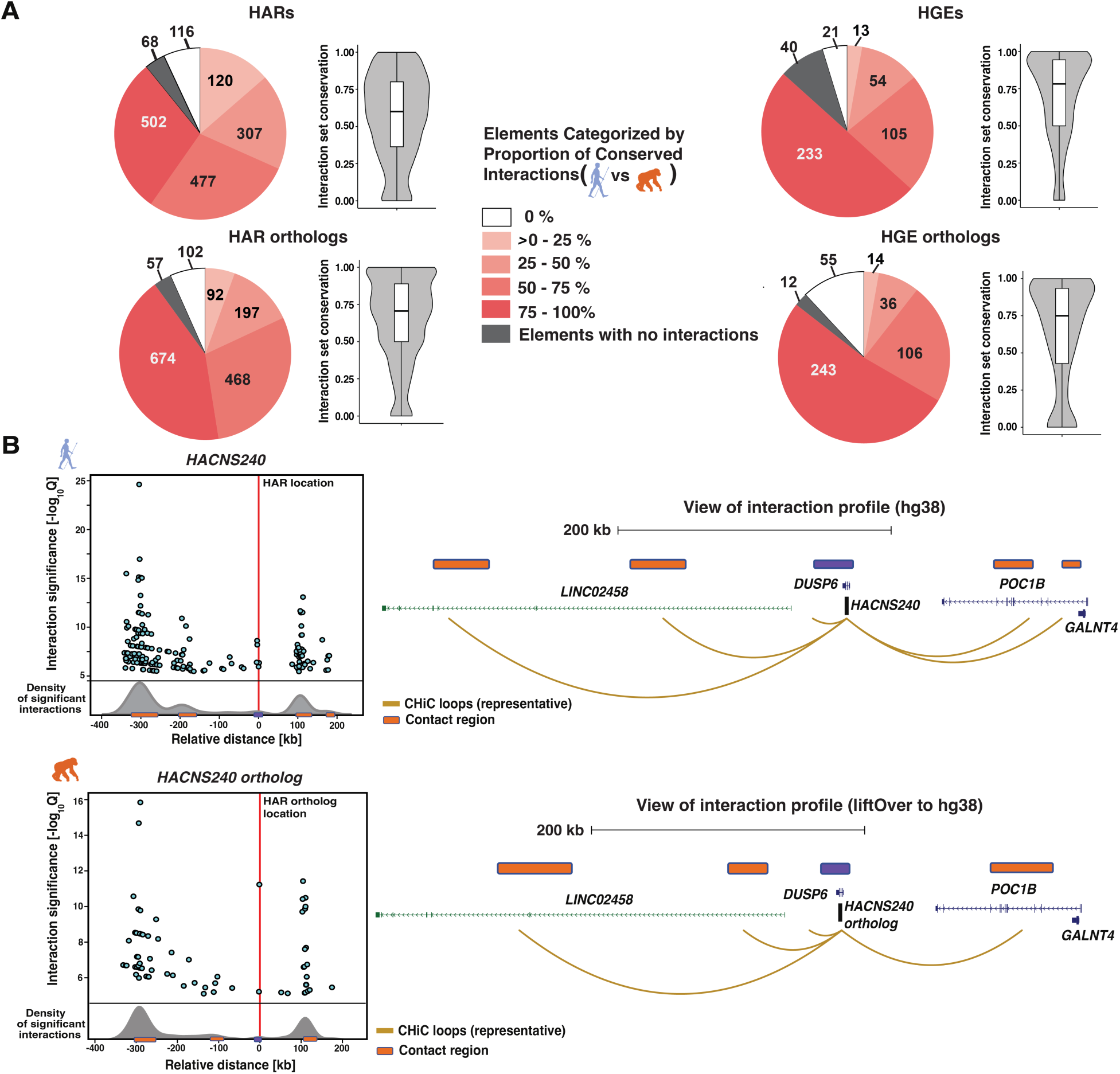
Conserved and species-specific HAR and HGE interactions in human and chimpanzee NSCs. **(A)** *Left.* Proportion of conserved interactions for HARs and their chimpanzee orthologs. The pie charts show the breakdown of HARs and chimpanzee orthologs by the percent of conserved interactions, in quartiles as shown in the legend. The violin plot shows the overall distribution of conserved interactions across HARs and their orthologs, and indicates the median value as well as the interquartile range. *Right*. Proportion of conserved interactions for HGEs and their orthologs, as shown for HARs on the left. **(B)** CHi-C interaction profiles of *HACNS240* (*top*) and its chimpanzee ortholog (*bottom*). *Left.* Each point on the scatterplot represents a significant interaction, plotted by the degree of significance as shown on the y-axis. The relative distance from the HAR is given on the x-axis. A normalized density distribution of the interactions is shown below the scatterplot, and regions of high interaction density are marked by orange boxes (Methods). *Right.* Schematized UCSC Genome Browser view (in GRCh38 coordinates) for each interaction profile. The location of *HACNS240* and its ortholog are shown in purple above the gene models, and the regions of high interaction density are shown in orange. Representative looping events are shown in gold. Protein-coding genes are shown in blue and lncRNA genes are shown in green.

To evaluate whether HARs and HGEs directly regulate their gene targets, we used both CRISPRi and CRISPRa to respectively decrease or increase the activity of one HAR, *HACNS52*, and its chimpanzee ortholog. This HAR is marked by H3K27ac in hNSCs, suggesting it is an active enhancer. *HACNS52* also contacts two genes, *ANXA2* and *ICE2* (Fig. S3A; Methods). Both of these target genes have been shown to have highly enriched expression in the progenitor zones of the human fetal brain^50,51^. The chimpanzee ortholog also interacts with the orthologous genes in cNSCs (Fig. S4A). We found that the expression of both gene targets decreased via CRISPRi-mediated repression of *HACNS52* and its chimpanzee ortholog, and expression increased upon CRISPRa-mediated activation (Fig. S3B-C, Fig. S4B-C). This result supports that the interactions between *HACNS52* and its gene targets are capable of conveying regulatory information from this HAR to its target genes.

### HAR and HGE conserved gene targets in NSCs converge on neurodevelopmental functions

Using the criteria described above, we found that 2,963 gene targets (68.4% of the set of gene targets in human and 72.9% of the set of gene targets in chimpanzee) were conserved between HARs or HGEs and their chimpanzee orthologs (Fig. 3A and Fig.S22; Table S1; Methods). Collectively, we identified gene targets for almost 89% of HARs, considerably more than previously reported (Table 1). Slightly more than half (51.5%) of these conserved gene targets were protein-coding genes and the remainder were annotated as long non-coding RNA (lncRNA) genes, most of which were not expressed in the bulk RNA-Seq datasets we generated from hNSCs or cNSCs (Fig. S5A; Methods). This suggests that HARs may not only regulate protein-coding genes but may also play a role in lncRNA-mediated gene regulation^52^, although potentially in other biological contexts than those we examined here. We next conducted a GO Biological Process (BP) enrichment analysis of the conserved protein-coding gene target set using EnrichR (Methods). We found that these gene targets were significantly enriched in neuronal functions including axon guidance, synaptic transmission, cell-cell adhesion and neuron migration (Fig. 3B; Table S1). This led us to hypothesize that HARs may be involved in interactions that are established but not active at the NSC stage, but which are activated upon neuronal differentiation. We address this question further in a subsequent section below.

**Figure 3.**
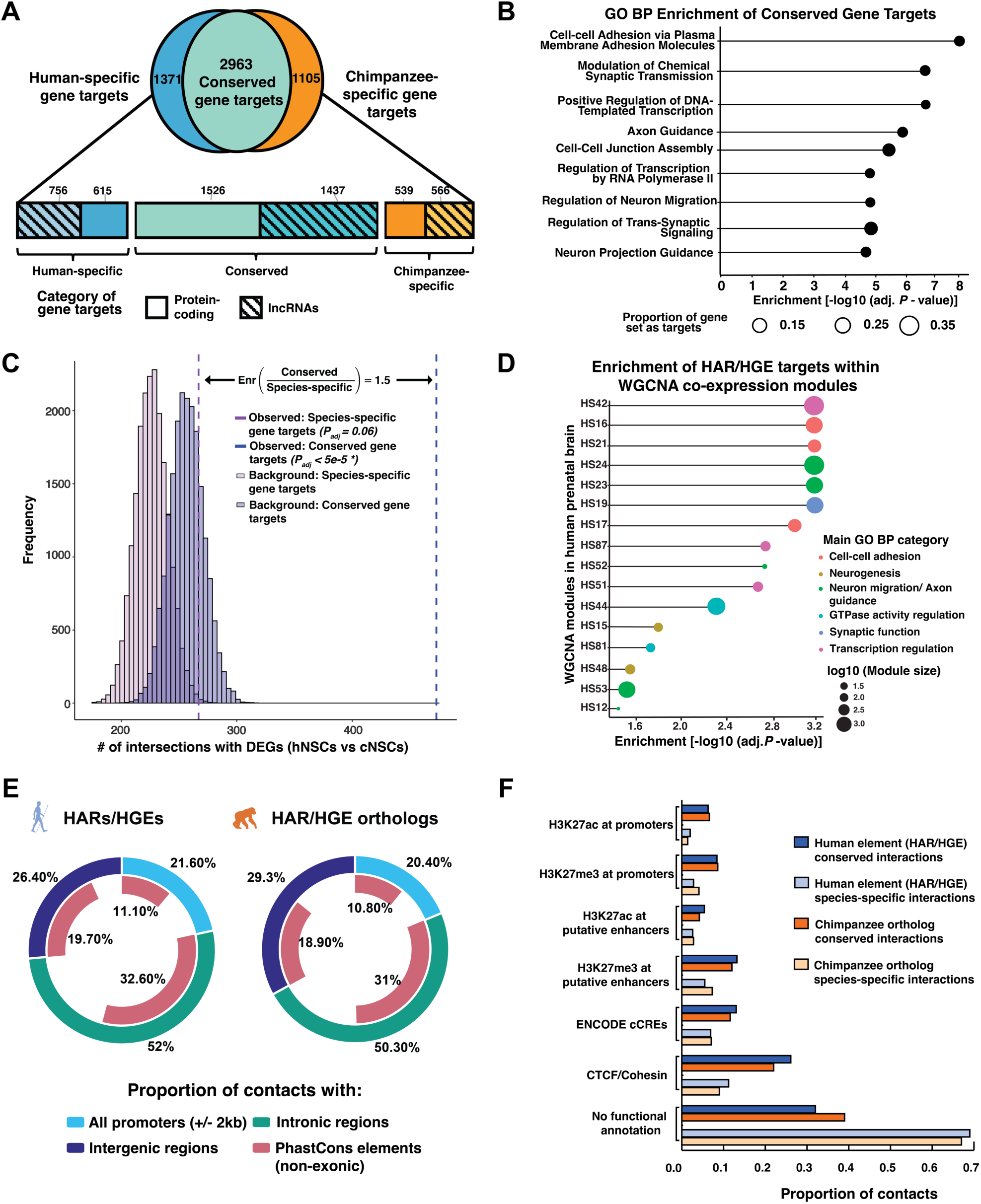
Functional annotation of HAR and HGE interactions in human and chimpanzee NSCs. **(A)** Conserved and species-specific gene targets of HARs, HGEs and their chimpanzee orthologs based on the human GENCODEv43 annotation, as defined in the Results and Methods. The horizontal bar plot further subdivides human-specific, chimpanzee-specific, and conserved targets into protein-coding and long non-coding RNA (lncRNA) genes. **(B)** GO Biological Process enrichment analysis performed on the conserved set of protein-coding gene targets, using the human GENCODEv43 annotation. **(C)** Enrichment of conserved HAR or HGE gene targets within differentially expressed gene (DEG) sets called between hNSCs and cNSCs and their relative over-representation within the DEG set compared to species-specific gene targets (Methods). *P*-values were computed based on permutation using random sampling of the background followed by Bonferroni correction (n = 20,000 trials; Methods). **(D)** HAR or HGE gene target enrichments in gene co-expression modules previously identified using WGCNA in human fetal brain. Modules start with the identifier ‘HS’ as they are called in human samples. The circles are colored according to significantly enriched GO Biological Processes as shown in the legend (Methods). The enrichment adjusted *P*-value for each module was calculated based on a permutation test with n = 20,000 trials followed by Benjamini-Hochberg (BH) correction. **(E)** Doughnut plot showing the distribution of HAR and HGE interactions based on gene and phastCons constrained noncoding element annotations as described in the Results and Methods. **(F)** Bar plot showing the proportion of conserved and species-specific interactions for HARs, HGEs and their chimpanzee orthologs overlapping putatively functional elements based on H3K27ac, H3K27me3, CTCF, and RAD21 profiles generated in this study and ENCODE cCREs.

It has been suggested that HARs may drive changes in gene expression by targeting different genes in human compared to their chimpanzee orthologs^7^. To investigate this mechanism, termed enhancer hijacking, we considered species-specific gene targets for each HAR and HGE. We initially identified 615 human-specific and 539 chimpanzee-specific protein-coding targets, based on whether an interaction with a gene was observed in one species but not the other using the criteria described above (Fig. S22, Methods). These gene sets were not enriched for any GO Biological Process (BP) category and did not show significant gene expression differences overall between human and chimpanzee NSCs (Fig S5B; Table S1). In contrast, we found that conserved HAR and HGE gene targets were significantly overrepresented among differentially expressed genes (DEGs) called between hNSCs and cNSCs while species-specific gene targets were not (Fig. 3C; Methods). Conserved gene targets were approximately 1.5 times more enriched among DEGs than species-specific gene targets. To determine if this result was influenced by the criteria we used to identify gene targets, we then defined a strict set of species-specific protein-coding gene targets as genes targeted in one genome with no interaction detected within a 50 kb window around the orthologous gene in the other genome (Fig. S22). This yielded 323 human-specific targets and 359 chimpanzee-specific targets. This refined target set was also not enriched for any GO BP category and did not show species differences in expression overall (Fig. S5C). These results suggest that species-specific HAR and HGE gene targets generally do not converge on any specific functions or show species-biased expression changes in NSCs.

It has also been suggested human-specific structural variants (hsSVs) may be associated with changes in HAR gene targeting between human and chimpanzee^7,53^. To explore this question, we first asked whether the HARs in our dataset were enriched in topologically associated domains (TADs) previously identified in the human fetal brain that also included hsSVs, as described in the previous study^25^. We found that HARs were enriched in TADs containing hsSVs compared to other evolutionarily constrained phastCons elements (Fig. S6A-C). This was also true for HARs at the extreme ends of the distribution of conserved interaction values shown in Fig. 2A. HARs with >90% of interactions conserved with their chimpanzee ortholog were enriched in TADs with hsSVs, as were HARs with <10% of their interactions conserved (Fig. S6D-E). We also observed similar enrichments for HGEs and transcriptional enhancers identified in transgenic mouse assays, which were defined based on constraint or histone modifications associated with enhancer activity (Fig. S6F-G; Methods)^22,54^. Collectively, these results support that HARs are not uniquely associated with hsSVs compared to other regulatory elements.

To gain further insight into the functional pathways in which HAR and HGE gene targets operate, we then asked whether gene targets were enriched in gene co-expression modules previously identified in the human fetal brain^34^. These modules are often enriched in genes that participate in common biological processes. We found that HAR and HGE gene targets are enriched in 15 out of 87 gene co-expression modules defined in the previous study (Benjamini-Hochberg (BH)-corrected *P* < 0.05; permutation test; Methods). Genes within these 15 co-expression modules were significantly enriched for neurodevelopmental functions, including cell-cell adhesion, neurogenesis, axon guidance and synaptic function (Fig. 3D). We also examined whether HAR and HGE gene targets were enriched in sets of genes associated with neurodevelopmental and neuropsychiatric disorder risk (Table S1). We found that HAR and HGE gene target sets were significantly enriched within gene sets associated with risk for ASD or schizophrenia (SCZ) (BH-corrected *P* = 0.04 for ASD, BH-corrected *P* < 5e-5 for SCZ; Fig. S7A-B; Methods)^55,56^. These findings further support that HARs and HGEs may play a role in regulating genes within neurodevelopmental pathways, as well as genes implicated in neurodevelopmental and neuropsychiatric disorders.

### HARs and HGEs interact with other putative gene regulatory elements

We next sought to identify interactions among HARs, HGEs and other regulatory elements in the genome. To accomplish this, we categorized interactions using two approaches. First, we used public annotations of constrained elements (identified via phastCons) and ENCODE candidate cis-regulatory elements (cCREs)^57,58^. Second, we used the maps of histone modifications and regulatory factor binding events we generated in human and chimpanzee NSCs to annotate interaction activity states. We found that approximately one-fifth of interactions were in promoter regions, defined within 2kb of an annotated transcription start site, while approximately half fell within introns and the remainder were in intergenic regions (Fig. 3E). Interactions were significantly enriched within introns (*P* < 5e-5, permutation test; Methods). This could be a consequence of dynamic loop extrusion during transcriptional elongation, in which a HAR or HGE interaction is captured at a particular location along an actively transcribed gene in a subset of cells,^59,60^ or interactions involving regulatory elements within introns^61^.

We then considered HAR or HGE interactions with non-exonic phastCons constrained elements, which include regulatory elements^62,63^. We found that phastCons elements were significantly enriched among HAR or HGE interactions: >60% of all interactions in human or chimpanzee involve a phastCons element (Fig 3E; *P<5e-5*, permutation test; Methods). We also found that 10% of all interactions in human and chimpanzee NSCs also involve ENCODE cCREs identified in human neural progenitors (Fig. 3D). Additionally, we identified a set of 22 HAR-to-HAR, 7 HAR-to-HGE and 2 HGE-to-HGE interactions, in which reciprocal interactions occur between genomic regions containing HARs or HGEs (Table S1; Methods). This indicates that HARs and HGEs interact with one another as well as with other regulatory elements in the genome.

To further explore potential functions of HAR or HGE interactions, we integrated the interaction maps with maps of histone modifications and binding events we generated as described above. We found that 56% of all HAR and HGE interaction partner loci (and 53% of the corresponding chimpanzee ortholog interaction partner loci) overlapped with at least one H3K27ac, H3K27me3, CTCF, or RAD21 peak or annotated ENCODE cCRE (Fig. S8B). We then compared conserved versus species-specific interactions. Both sets of interactions showed similar overlap with phastCons elements (average overlap = 58.3% for the conserved set, average overlap = 56.8% for the species-specific set; Fig. S8A). However, a lower proportion of species-specific interactions in both human and chimpanzee overlapped with one of the five features described above compared to conserved interactions (Fig. 3F). Overall, species-specific interactions were significantly less frequently associated with any of these features than conserved interactions (Fig. 3F; *P < 0.05,* Fisher’s exact test; Methods). Collectively, these results suggest that species-specific interactions are less likely to involve regulatory elements relevant to NSC biology or may involve functions not captured by our datasets.

### Linking HARs and HGEs to species differences in gene target expression

We next sought to link potential differences in HAR and HGE regulatory activity between human and chimpanzee with changes in gene expression. We first considered HARs and HGEs that were marked by H3K27ac in either cNSCs or hNSCs and compared changes in H3K27ac levels at each HAR or HGE with changes in gene target expression (Fig. 4A and Fig. S9A). We found a positive correlation between the magnitude of changes in H3K27ac levels at HARs or HGEs and the magnitude of target gene expression changes (Pearson correlation π = 0.36; *P <* 2.2e-16; the curve of best fit is a cubic polynomial based on minimizing residual squared error; Methods). This supports a general trend between species differences in H3K27ac levels at HARs or HGEs and the species bias in the expression of gene targets. When we considered conserved and species-specific targets separately (Fig. S9B), we found that conserved target gene expression exhibited a higher correlation with H3K27ac levels (π = 0.38) compared to the species-specific targets (π = 0.27).

**Figure 4.**
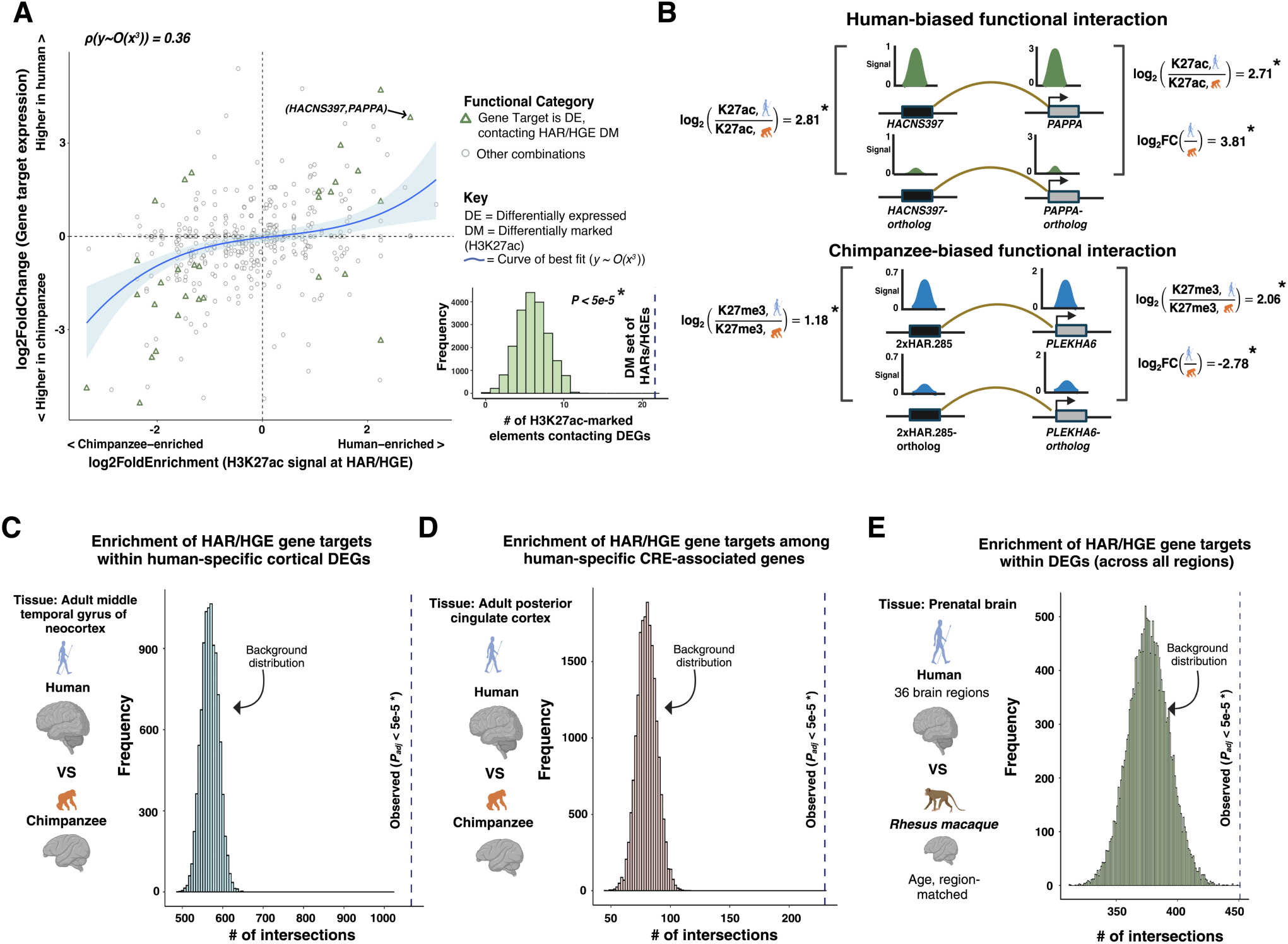
HARs and HGEs are significantly associated with gene expression differences between human and chimpanzee. **(A)** Scatterplot of relative expression for all HAR and HGE gene targets (log2(human TPM/chimpanzee TPM)) against the relative H3K27ac level at the HAR or HGE (log2(human/chimpanzee)). Differentially expressed (DE) gene targets contacted by elements showing differential H3K27ac marking (DM) are plotted as green triangles. The polynomial of best fit (y ∼ O(x^3^)) is plotted in blue and standard error is shaded in light blue; see Results and Methods for details. The inset shows the expected distribution of HARs and HGEs differentially marked by H3K27ac that also target a differentially expressed gene as obtained by permutation, compared to the observed value shown by the vertical dotted line. The enrichment *P*-value was computed based on random sampling of the background (n = 20,000 trials; Methods), * = *P* < 0.05. **(B)** Two examples of HAR gene targets whose species-biased differential expression is associated with a specific enrichment of H3K27ac (in green) or H3K27me3 (in blue) at the HAR and the target gene promoters as described in the Results. Fold enrichment and fold change (FC) ratios are shown on a log2 scale, and significant differences in marking and expression were identified using DESeq2 (* = BH-corrected *P* < 0.05). **(C – D)** Enrichment of HAR and HGE gene target sets within **(C)** human-specific DE genes and **(D)** human-specific differentially accessible region (CRE)-linked genes, respectively, as described in the Results. *P*-values were computed based on permutation using random sampling of the background (n = 20,000 trials; Methods), * = Bonferroni-corrected *P* < 0.05. The observed intersection of these gene sets with HAR and HGE targets in hNSCs is shown by the dotted line. **(E)** Enrichment of HAR and HGE gene target sets within differentially expressed genes identified in a comparison of human and rhesus macaque fetal brain as described in the Results. The DEGs were identified as differentially expressed in at least one brain region. *P*-values were computed as in **(C – D)** with * = Bonferroni-corrected *P* < 0.05, and display of observed intersection with hNSC gene targets is shown by the dotted line as in (**C – D)**. HAR and HGE targets were combined for these analyses. Illustrations in these panels were generated using BioRender.com.

We then sought to identify individual examples of changes in HAR or HGE activity that were linked to changes in gene target expression. We used DESeq2 to identify HARs and HGEs that showed significant differences in H3K27ac levels between hNSCs and cNSCs, as well as target genes that showed significant differences in expression (DEGs; Methods). We found that differentially marked HARs and HGEs were significantly more likely to target a DEG compared to HARs that did not show significant differences in marking between human and chimpanzee (enrichment ratio = 2.64, *P* < 5e-5, permutation test; Fig. 4A; Methods). This supports that changes in enhancer activity at particular HARs and HGEs are associated with changes in expression of particular gene targets between both species. We identified 21 HARs or HGEs that were significantly differentially marked and that also targeted a DEG (Fig. 4A; TableS2). One example is shown in Fig. 4B (top) and Fig. S10. In this case, *HACNS397* exhibited significantly greater levels of H3K27ac in hNSCs and targeted *PAPPA*, whose promoter showed a significant increase in H3K27ac. *PAPPA* also showed significantly increased expression in hNSCs compared to cNSCs (Fig 4B, *top*). *PAPPA* encodes a secreted metalloproteinase expressed during early human brain development and has been shown to play a role in regulating the rate of embryonic development in zebrafish models^64,65^.

In contrast to the positive correlation between changes in the level of H3K27ac marking and gene expression we observed, we detected a weak negative correlation between changes in H3K27me3 levels and gene expression changes (Pearson correlation ρ = -0.16 for curve of best fit; Fig. S11A). However, we did identify 11 HARs or HGEs exhibiting significant changes in H3K27me3 levels that also targeted DEGs (Table S2). For example, 2xHAR.285 showed significantly increased levels of H3K27me3 in hNSCs compared to its ortholog in cNSCs and targets *PLEKHA6*, whose promoter also showed increased H3K27me3 (Fig 4b, *bottom*; Fig. S11B). *PLEKHA6* is a chimpanzee-biased DEG showing significantly greater expression in cNSCs compared to hNSCs (Table S2), and encodes an adherens junction protein that regulates copper homeostasis in the brain^66,67^.

We then examined whether HAR targets included previously identified genes with human-specific expression changes in the developing and adult brain^32–34^. We initially considered two studies comparing human and chimpanzee adult brain regions (Figs. 4C, 4D). The first study identified genes with significantly increased expression in human using scRNA-seq data, ^32^ and found that genes located near HARs in two-dimensional, linear genomic terms were enriched among these human-biased genes. A second study using scATAC-Seq found that HARs were enriched in human-specific differentially accessible regions in the brain, many of which had nearby genes showing species-biased expression^33^. However, these studies relied on assigning potential gene targets to HARs based either on their linear proximity in the genome, or by assigning HARs to putative regulatory regions and then linking them to nearby genes, without any direct evidence of interaction. Using our set of empirically defined HAR and HGE gene targets in hNSCs, we found that genes directly targeted by HARs and HGEs were highly enriched among human-specific DEGs and human-specific differentially accessible region-linked genes showing biased expression in adult brain regions (Figs. 4C and 4D; *P_BH_ < 5e-5*, permutation test). We then considered a study comparing gene expression in multiple fetal brain regions in human and rhesus macaque^34^ (Fig. 4E, *P < 5e-5*, permutation test). HAR and HGE gene targets were enriched among the union of all DEGs identified in the study. However, gene targets were not enriched within DEGs identified in comparisons of specific brain regions, such as human versus macaque pre-frontal cortex (Fig. S14B). As observed for DEGs in hNSCs and cNSCs (Fig. 3C), we found that conserved gene targets were more highly enriched than species-specific targets among sets of genes identified from each of these human-chimpanzee and human-macaque brain comparisons, as well as among DEGs called between human and chimpanzee hippocampal intermediate progenitors cultured from iPSC lines (Fig. S12A-B, Fig. S13A-B)^68^. Gene targets were not enriched in DEGs identified in non-neuronal tissues, specifically kidney and liver, supporting that the enrichments we observed may be specific to the brain (Fig. S14C-D)^69^. GO BP enrichment analysis on the set of gene targets present in each of the sets of DEGs or genes associated with differentially accessible regions identified in the previous studies highlighted a strong overrepresentation of neurogenic processes, such as cell-cell adhesion, ion-transport and synaptic transmission (Fig. S15A-F). Gene targets represented across all three DEG datasets included *CAMKMT*, which encodes a methyltransferase involved in the regulation of ion channels and *CUX1*, a transcription factor that plays a critical role in the formation of dendritic spines and synapses in layer 2-3 excitatory neurons of the cerebral cortex^70,71^.

### HARs and HGEs participate in active and repressed regulatory interactions in NSCs that change chromatin state upon neuronal differentiation

Our finding that HAR and HGE gene targets identified in hNSCs are involved in neuronal processes suggests that some HAR/HGE-gene target interactions may be silent or repressed in hNSCs but active in neurons. To address this question, we considered whether HAR/HGE-gene target interactions identified in hNSCs show changes in chromatin states and gene expression upon neuronal differentiation. We first compared HAR and HGE interaction maps and H3K27ac and H3K27me3 marking between hNSCs and iPSC-derived human neurons. Overall, we identified sets of shared as well as cell type-specific gene targets for HARs or HGEs (Fig. S16A). GO Biological Process (BP) enrichment analysis showed that gene targets shared between both cell types were significantly enriched in neuronal functions, such as cell-cell adhesion and synaptic processes, while the hNSC-specific protein-coding targets were enriched for progenitor-related functions, such as FGF signaling (Fig. S16B-C)^72–74^. However, GO analysis of the 492 human neuron-specific protein-coding gene target set showed no significant enrichment. We note that gene targets shared between both cell types but annotated as encoding neuronal functions may also have functions in NSCs that have yet to be determined.

We next considered the 533 HAR/HGE-gene target interactions that were marked by H3K27me3 at both interacting loci in hNSCs. We found that 20 interactions showed a switch from H3K27me3 to H3K27ac at both the HAR or HGE and the target genes (Fig. 5A; Table S2). One example, HAR116 and its gene target *TCF20*, is shown in Figs. 5B and 5C. Corresponding with the observed change in chromatin state, *TCF20* was not expressed in hNSCs but was expressed in human neurons (Fig. 5C). Activation of *TCF20* is consistent with its role in neuronal differentiation from later stage NSCs to neurons during cortical neurogenesis^75,76^. However, the majority of H3K27me3-marked interactions in hNSCs either maintained H3K27me3 in human neurons or lost H3K27me3 marking without a corresponding gain of H3K27ac (Fig. 5A). For example, the interaction between *HACNS224* and *CDH23* maintained H3K27me3 marking and correspondingly low levels of *CDH23* expression in both cell types (Fig. 5C, Fig. 5D), while the interaction between *HACNS475* and *NKX6-2* and *CFAP46* lost H3K27me3 marking at the HAR during neuronal differentiation (Fig. S18A-B). To evaluate whether maintenance or conversion of repressed and active chromatin states between hNSCs and neurons corresponded to overall changes in gene target expression, we compared the expression distributions of gene targets for each of the three interaction categories shown in Fig. 5A. Gene target expression did not significantly increase when H3K27me3 marking was maintained in both hNSCs and neurons or lost without subsequent gain of H3K27ac (Fig. 5E). However, a switch from H3K27me3 marking in hNSCs to H3K27ac in neurons was associated with significantly increased gene target expression (BH-corrected *P* = 0.004; median log2(neuron TPM/hNSC TPM) = 3.4). This supports that a switch from a repressed to an active chromatin state at HARs or HGEs and their gene targets upon neuronal differentiation is associated with upregulation of gene target transcription.

**Figure 5:**
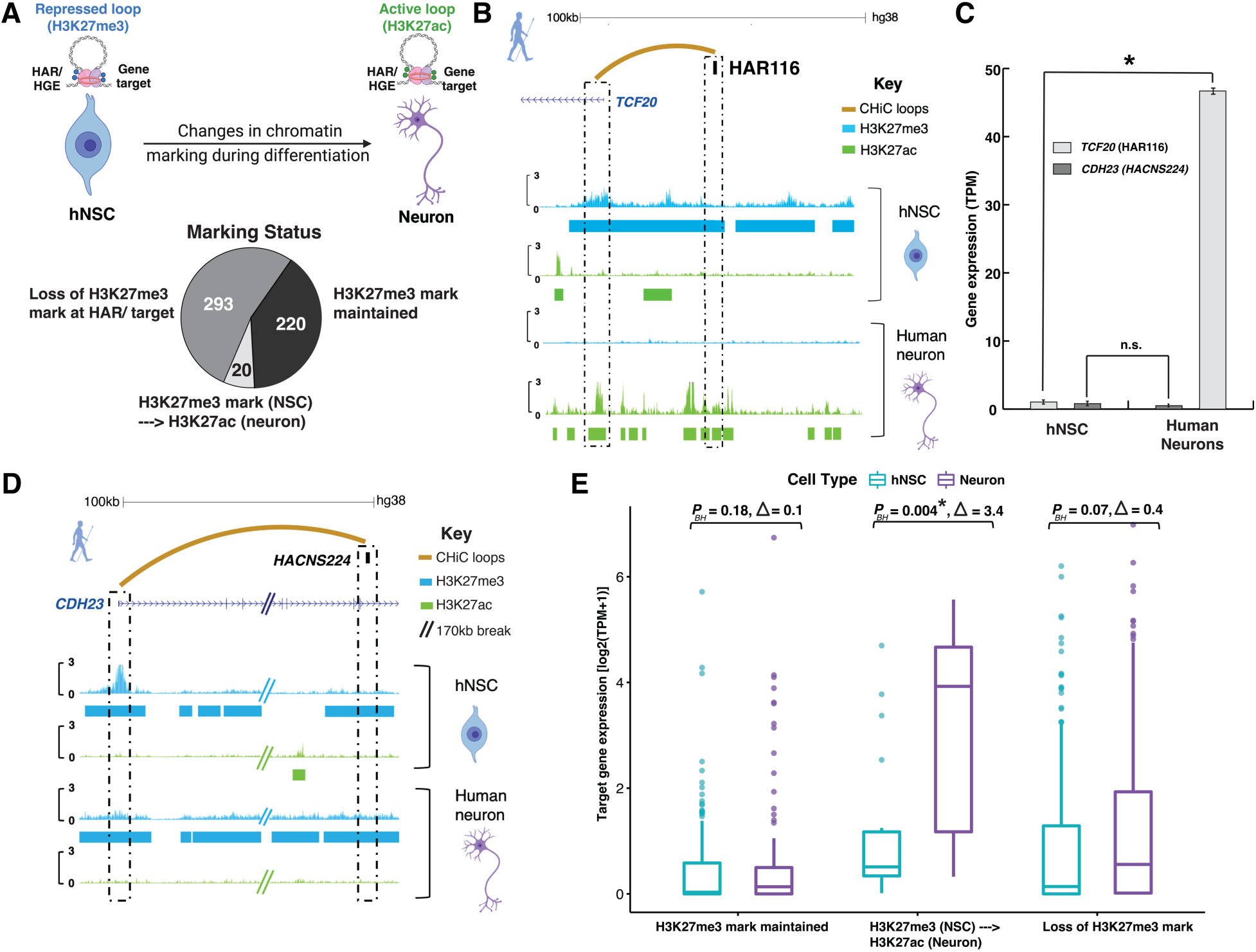
Activation of repressed HAR and HGE interactions upon neuronal differentiation. **(A)** The number of H3K27me3-marked interactions in hNSCs that maintain H3K27me3 marking in human neurons (black), interactions that switch to an H3K27ac-marked state in neurons (white) and interactions that lose H3K7me3 at either the HAR, HGE or the gene target (grey). **(B)** An example of a repressed interaction that is activated upon neuronal differentiation. HAR116 engages in an H3K27me3-marked interaction with its gene target *TCF20* in hNSCs, which switches to an H3K27ac-marked state in neurons. The curved golden line indicates the HAR116-*TCF20* interaction detected via CHi-C, and H3K27me3 (in blue) and H3K27ac (in green) profiles are shown for both hNSCs and iPSC-derived human neurons. Dashed black boxes highlight the location of the HAR and the *TCF20* promoter relative to the H3K27me3 and H3K27ac peaks. **(C)** Comparison of of *TCF20* and *CDH23* expression in hNSCs and neurons (TPM values, P_BH_ < 0.05 by Mann-Whitney test). **(D)** Example of a repressed interaction in NSCs that is maintained in neurons, shown as in **(B)**. **(E)** Whisker plots with pairwise comparisons of distributions of gene target expression for each category of H3K27me3-marked interactions across hNSCs (in teal) and neurons (in purple). Each box represents the 25^th^ to 75^th^ percentiles of the data, and the median of each distribution is marked with a solid line. The median increase in log2(TPM+1) between neurons and hNSCs for each of the three categories is denoted by Δ. *P* values for differences in expression between NSCs and neurons in each category were calculated using a Mann-Whitney U test (* = BH-corrected *P* < 0.05).

### HAR and HGE gene targets show cell type-specific expression patterns in the fetal human brain

We next leveraged the gene target sets we identified to investigate the cell types in which HARs and HGEs may have altered gene expression in the developing human brain. To accomplish this, we integrated gene target sets from hNSCs or human neurons with a previously generated atlas of single-cell gene expression in the fetal human brain, spanning Carnegie stage 14 (CS14) to gestational week 25 (GW25)^35,36^.We analyzed expression in seven regions of the developing forebrain and the cerebellum, and identified progenitor and neuronal cell types based on previously defined sets of marker genes. We then quantified the average expression of hNSC and human neuron gene targets in 29 progenitor and 33 neuronal subtypes, respectively.

We first sought to identify whether the expression profiles of HAR and HGE gene targets identified in hNSCs converged on specific progenitor populations in the developing human brain. We partitioned hNSC gene target sets into 15 clusters using k-means clustering on their average expression profiles across progenitors in the eight brain regions included in the atlas (Methods). Clusters of gene targets exhibited brain region-specific expression profiles, including a cluster of 21 genes with cerebellum-biased expression and 42 genes with expression biased towards ventral forebrain regions (Fig. S19). Multiple clusters of gene targets also exhibited an expression bias towards specific cortical progenitor subtypes, including *HOPX-*positive outer radial glia (oRG, 52 gene targets contacted by 86 HARs and HGEs) and *EOMES*-positive intermediate progenitor cells (IPCs, 38 gene targets contacted by 63 HARs and HGEs; Fig. 6A). Both oRGs and IPCs have been implicated in primate-specific cortical expansion^77–80^. These clusters also showed patterns of higher expression in *EOMES*-positive and *HOPX*-positive progenitor subtypes of other sampled brain regions including the hippocampus, hypothalamus, and cerebellum (labeled as NSC2 and NSC3 in Fig 6A). The oRG-biased group included multiple gene targets with known neurodevelopmental functions including *ANXA2* and *ROBO1*^81–83^. We also identified multiple IPC-biased gene targets that have characterized neurodevelopmental phenotypes including *RBFOX2, AUTS2* and another ROBO family paralog, *ROBO2* (Fig. 6B)^84–86^.

**Figure 6.**
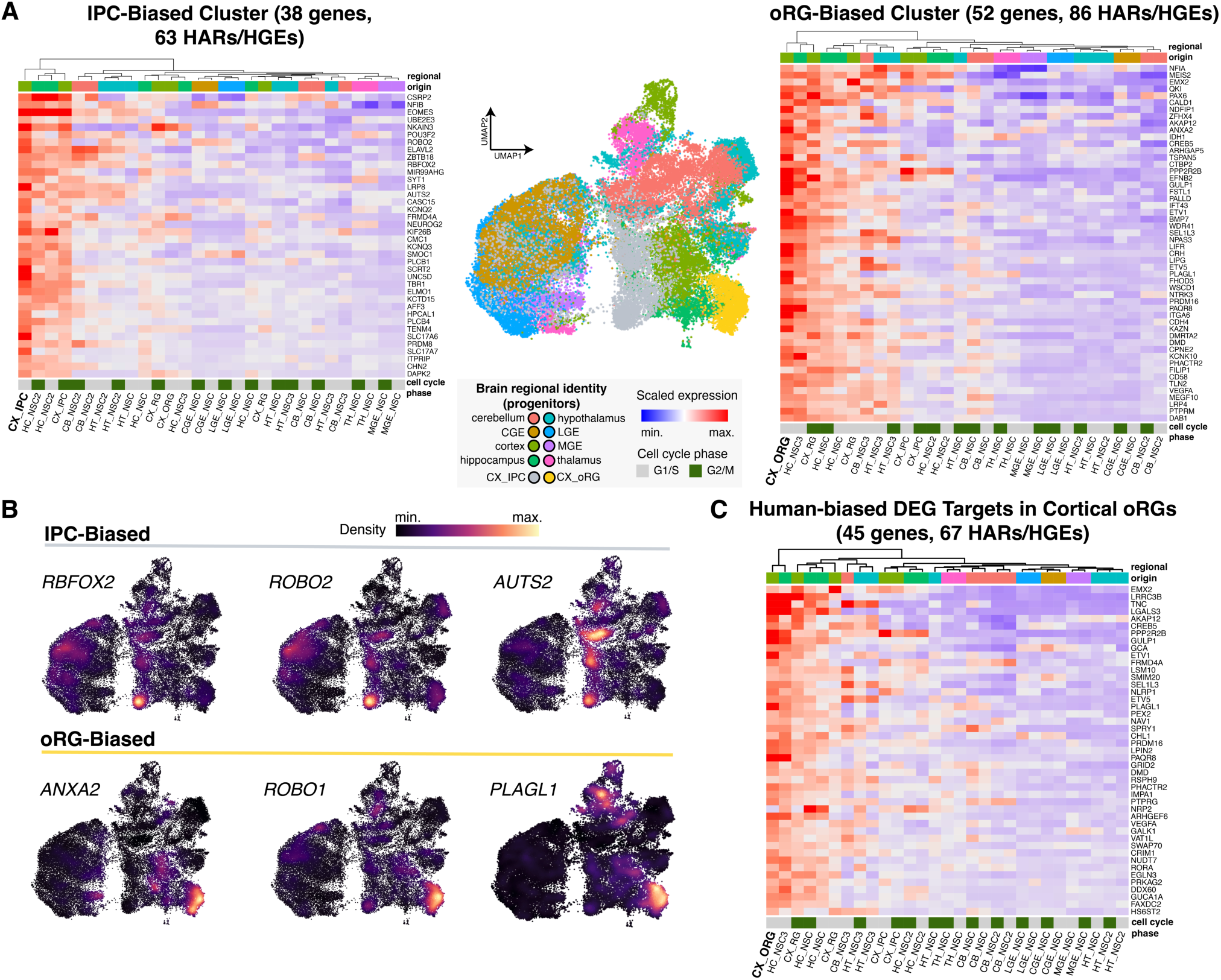
HAR and HGE gene targets in NSCs show cell type-specific expression profiles in the fetal human brain. **(A)** UMAP showing neural progenitors in scRNA-seq from eight embryonic and fetal human brain regions colored by regional identity with cortical outer radial glia (oRGs) and intermediate progenitor cells (IPCs) highlighted in yellow and silver, respectively. Heatmaps of clustered average scaled expression profiles across 29 neural progenitor subtypes, separated into gene sets that show cell-type specific expression in cortical oRGs and IPCs. CX – cortex; HC – hippocampus; HT – hypothalamus; CB – cerebellum; TH – thalamus; LGE – lateral ganglionic eminence; MGE – medial ganglionic eminence; and CGE – caudal ganglionic eminence. **(B)** Density plots for a subset of genes showing cell-type specific expression in progenitor cell types, visualized on the UMAP shown in A. **(C)** Visualization of the average expression profiles of 38 genes showing higher expression in hNSCs compared to cNSCs that share oRG-biased expression profiles.

Following on these results, we then investigated whether HAR and HGE gene targets that showed significantly greater expression in hNSCs compared to cNSCs showed cell-type specific expression biases. We identified a set of 45 DEG targets with human-biased expression in cortical oRGs and *HOPX*-positive NSCs in the hippocampus (Fig. 6C). These genes included *PLAGL1*, a transcription factor shown to promote neurogenesis in mouse neocortical neural stem cells *in vivo*, *TNC*, a glycoprotein component of the extracellular matrix that restricts differentiation and maintains stemness of spinal cord neural stem cells *in vitro*, and *PRDM16*, a zinc finger transcription factor and histone methyltransferase involved in fetal and postnatal neural stem cell maintenance, laminar specification of upper layer cortical neurons, and cortical neuron positioning ^77,87–90^.

We then asked whether HAR and HGE gene targets identified in human neurons also converged on certain brain regions or neuronal subtypes. We quantified and clustered the average expression profiles of all neuronal HAR and HGE targets in various neuronal subtypes of the eight profiled brain regions as described above. We identified gene target clusters that exhibited brain region-specific expression profiles, including in the thalamus, hypothalamus, or hippocampus, or in the hippocampus and cortex (Fig. 7A). Multiple clusters also showed biased expression in individual neuronal cell types, including rostral thalamic neurons (RTN), identified based on the coexpression of *SOX14* and *GATA3*^91^, and a population of hypothalamic glutamatergic neurons expressing *IRX1* and *IRX2* (Fig. S20). Specific examples of gene targets expressed within these neuronal cell types are shown in Fig. 7B. Gene targets with biased expression in RTNs include *CDH8*, a type II cadherin involved in the regulation of synaptic transmission across multiple brain regions, and *FOXP2*, a vital neurodevelopmental transcription factor that is enriched in primate-specific parvocellular neurons of the dorsolateral geniculate nucleus and is otherwise responsible for the specification of posterior thalamic nucleus identity ^92–96^. Additionally, *RBFOX1*, a neuron-specific splicing factor known to regulate the alternative splicing of neurodevelopmental genes, is a gene target present in the hippocampus- and cortex-biased cluster^97,98^. Overall, these findings provide a basis for hypothesis-directed studies examining how HARs and HGEs may have contributed to human-specific expression changes in a cell type-specific manner, and in multiple regions of the developing brain.

**Figure 7:**
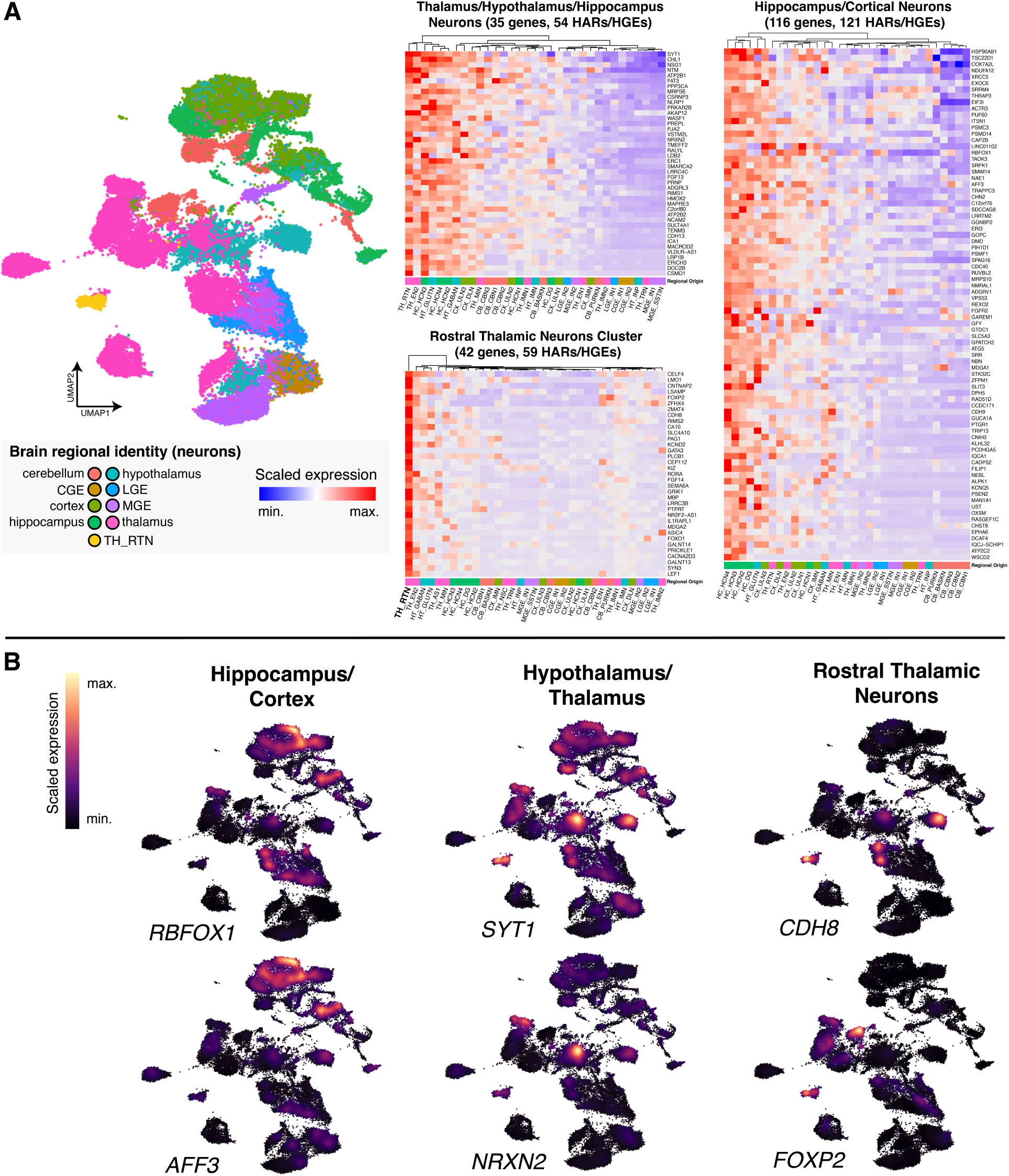
HAR and HGE gene targets in neurons show biased expression across multiple neuronal subtypes in the developing human brain. **(A)** UMAP showing neurons in scRNA-seq from eight embryonic and fetal human brain regions colored by regional identity with rostral thalamic neurons (RTN) highlighted in yellow. Heatmaps of clustered average scaled expression profiles across 33 neuronal subtypes, separated into gene sets with biased expression in groups of brain regions (thalamus/hypothalamus/hippocampus and hippocampus/cortex) and specific neuronal subtypes (rostral thalamic neurons). CX – cortex; HC – hippocampus; HT – hypothalamus; CB – cerebellum; TH – thalamus, LGE – lateral ganglionic eminence; MGE – medial ganglionic eminence; and CGE – caudal ganglionic eminence. The neuron subtype abbreviations are provided in Table S3. **(B)** Density plots of expression in neuronal cell types (visualized on the UMAP shown in A) display the regional or cell type-specific biases of HAR and HGE gene target expression within each cluster.

## Discussion

Understanding the role of HARs and HGEs in the evolution of the human brain requires the systematic identification of the genes they regulate during neurodevelopment and the biological pathways and cell types in which those genes function. Here we generated a high-resolution map of gene and gene regulatory element interactions involving HARs and HGEs in neural stem cells and neurons. We identified gene targets for almost 90% of HARs, many of which converge on neurodevelopmental processes including axon guidance, neuron migration and synaptic transmission, and which show regional and cell type-specific expression patterns in the human fetal brain. Our findings also support that HARs and HGEs may have contributed to the evolution of uniquely human brain features by altering the expression of a conserved set of genes targeted in both human and chimpanzee. Collectively, our results provide insight into a general mechanism by which HARs and HGEs drove changes in gene expression and will inform hypothesis-directed experimental studies of HARs and HGEs.

Our findings will significantly advance *in vivo* analyses of HARs and HGEs. Determining the specific traits that HARs and HGEs influenced ultimately requires genetic models, such as humanized mice or cerebral organoids, in which their effects on neurodevelopment and function can be measured. A small number of HARs and HGEs have been shown to drive changes in gene expression and regulation in humanized mouse models, as well as developmental phenotypes including changes in radial glial self-renewal that lead to an increased number of neurons in the cortex^12,15,24^. These studies all share two advantages: first, strong evidence that the HAR or HGE encoded human-specific regulatory activity in a particular tissue, developmental time point or cell type; and second, a clear understanding of the identity and biological function of the likely gene targets. This enabled a hypothesis-directed, targeted analytical approach focused on characterizing phenotypes in the biological contexts where they were most likely to emerge. However, to date we have lacked this level of prior knowledge for the vast majority of HARs. High-throughput approaches, such as MPRAs or comparisons of biochemical signatures associated with enhancer activity across species, as well as low-throughput transgenic enhancer assays, have identified HARs and HGEs that exhibit human-specific changes in regulatory function. Our findings now make it possible to link these HARs and HGEs to genes with known biological functions in specific cell types, particularly to genes with prior evidence of expression changes in human. This will facilitate hypothesis-directed studies of many more HARs and HGEs using genetic models. Moreover, our results provide a basis for combinatorial modeling of HARs and HGEs that regulate genes in a common pathway or cell type, such as oRGs. Although previous *in vivo* studies of individual HARs and HGEs have identified clear phenotypes, the overall magnitudes of those phenotypes have been modest. Mouse or organoid models incorporating multiple HARs or HGEs with gene targets with known convergent functions and human-specific changes in their expression would allow us to understand how HARs or HGEs acted to rewire neurodevelopmental pathways and reveal complex phenotypes that recapitulate uniquely human brain features.

Our study also addresses a major question in the field: whether HARs acted by altering the expression of a common set of genes targeted by HARs and their orthologs in human and chimpanzee, or by HARs gaining new gene targets specifically in human via enhancer hijacking. Our results support the first mechanism. The majority of HAR and HGE gene targets were conserved between human and chimpanzee, converged on known neurodevelopmental pathways, and were overrepresented among differentially expressed genes identified in our study and in multiple prior studies. Species-specific targets did not show similar signatures, supporting that enhancer hijacking is not the primary mechanism by which HARs function. We propose that this finding is consistent with a conservative model of human brain evolution, in which unique features of the human brain arose via modification of otherwise highly conserved developmental processes and pathways. However, we acknowledge that individual species-specific changes in gene targets may have contributed to human-specific changes in brain development and function. It is also possible that enhancer hijacking events involving other classes of regulatory elements with human-specific activity, such as hCONDELs or HAQERs, may have influenced the evolution of human traits, or that HARs may engage in enhancer hijacking in biological systems we did not study here^99–101^. Related to this question, HARs were not specifically associated with human-specific structural variants (hsSVs) compared to HGEs or known enhancers in our analysis. A role for hsSVs in HAR evolution therefore remains to be established.

We also identified an additional feature of HAR and HGE regulatory function: that some interactions are established in a repressed state in NSCs and then switch to an active state in neurons. This suggests some HARs and HGEs, such as HAR116, are involved in neuronal fate determination, particularly for pro-neurogenic genes such as *TCF20*^75,76^. Such a mode of action may implicate these HARs and HGEs in the formation of “permissive” interactions, where interactions may be pre-set between the HAR or HGE and the gene target at the progenitor state and then become activated upon neuronal differentiation, as previously observed for enhancers in *Drosophila* and mouse^31,102^. More detailed work is needed to determine the biological significance of these cases and any role they may play in the evolution of uniquely human brain features.

By integrating our list of HAR and HGE gene targets with public human embryonic and fetal brain single cell expression atlases, we were able to identify the cell types in which these targets were likely to function during human brain development. This provides a basis for assigning HARs and HGEs to specific cell types, opening up a new avenue of investigation to study the roles of HARs and HGEs in the development, function and potentially the evolution of those cell types in human. For example, our findings implicate multiple HARs and HGEs in outer radial glia (oRG) biology. This provides an opportunity for studying the role of these HARs and HGEs in oRG specification and proliferation, and potentially in human cortical expansion. Notably, our results identified HAR and HGEs that may regulate genes such as *TNC*, known for their role in oRG development in the brain^77^. We also linked HARs and HGEs to other genes implicated in human brain evolution, including *ZBTB16*, known for its role in the generation of upper layer cortical neurons and *ZEB2*, which has been implicated in the evolutionary expansion of the human forebrain^103,104^. We note that our results do not directly demonstrate that HARs or HGEs regulate these or other genes in specific cell types in the developing brain. However, they provide a basis for hypothesis-directed experiments to study HAR and HGE-driven regulatory changes in specific cell types in model systems. The ongoing generation of single-cell expression data and other functional data in developing and adult human and non-human primate brain, including higher-resolution and potentially cell-type specific chromatin interaction data, will further inform these studies.

Our study has several limitations. We generated HAR and HGE interaction maps in cultured neural stem cells and neurons. Extending our approach to organoid models and to developing and adult brain tissues would provide additional insight. Moreover, by focusing solely on HARs and HGEs, we were not able to globally characterize regulatory interactions for all the gene targets we identified. As many genes are regulated by multiple enhancers, understanding how HARs and HGEs function within these larger, complex regulatory architectures will require additional analyses. These include comprehensive maps of regulatory interactions for all gene targets, as well as high-throughput perturbation of HARs, HGEs, gene targets and other regulatory elements to determine effects on gene expression and potentially identify cellular phenotypes. We also note that we have not interrogated HAR or HGE interactions in other developmental contexts, such as the limb, in which HARs and HGEs may have contributed to the evolution of uniquely human traits^10,105^. Still, our approach provides a framework for these studies, and for the comprehensive analysis of the thousands of loci in the genome identified to date that may encode human-specific regulatory functions underlying the unique biological features of our species.

## Acknowledgements

We thank S. Mane, C. Castaldi and E. Sykes at the Yale Center for Genome Analysis for generating the sequencing data used in this study; Y. Gilad, I. Gallego Romero and colleagues for providing human and chimpanzee iPSC lines; A. Bhaduri for assistance with accessing and interpreting the human fetal brain single-cell transcriptome data; G. Ganguly for help with the overall layout and aesthetics of figures and supplementary figures; and the members of the Noonan laboratory for their input on the manuscript. This work was supported by an award from the Eunice Kennedy Shriver National Institute of Child Health and Human Development (NICHD; R01 HD102030, to J.P.N); NSF Graduate Research Fellowships (to M.A.N., M.M. and K.M.Y.); an NICHD F32 Postdoctoral Fellowship (F32 HD108935, to M.B); and a Research Fellowship (352711928) from the Deutsche Forschungsgemeinschaft (DFG) (to S.U.). This research program and related results were also made possible by the support of the NOMIS foundation (to J.P.N.).

## Author Contributions

A.P., S.U., and J.P.N. conceived and designed the study. A.P. designed and performed CHi-C, CUT&RUN, RNA-Seq and CRISPR validation experiments. A.P. and M.A.N. cultured hNSCs and cNSCs. A.P. performed the differentiation of hNSCs to neurons. A.P., M.M. and R.P. conducted computational analyses. M.A.N. optimized NSC differentiation protocols and performed cell culture, bulk RNA-Seq, immunofluorescence imaging, marker gene qRT-PCR, and multiplexed scRNA-seq on hNSCs and cNSCs. M.M. designed and carried out the analysis of the fetal brain single cell expression data. J.W.Y. performed RNA extraction from human neurons prior to library preparation. S.U., M.B. and K.M.Y. provided advice on statistical approaches and visualizations. A.P. and J.P.N. wrote the paper with input from all authors.

## Data and code availability

All data generated in this study is available at the Gene Expression Omnibus under accession number GSE270272.

All original code generated for this study is available at GitHub: https://github.com/Noonan-Lab/Pal_et_al_HAR_interactome, https://github.com/NoonanLab/XSAnno_runs and Zenodo:10.5281/zenodo.12110226, 10.5281/zenodo.12521205.

## Methods

### Cell lines

All cell lines used in the study are listed in Table S1. Human and chimpanzee iPSC cell lines were provided by the Gilad laboratory and were generated as described^106^.

### Neural stem cell differentiation from iPSCs

Human and chimpanzee iPSCs were differentiated into neural stem cells using dual SMAD inhibition (STEMDiff. SMADi neural induction kit; catalog #08581). Briefly, iPSCs were plated at high density (2 million cells/well in a 6 well plate) on hESC-qualified Geltrex (Gibco #A1413302), then cultured in neural induction media at high density with ROCK inhibitor Y-27632 (StemCell Tech, catalog #72304). Cells were maintained at high density for 3 days before passaging every 48 hours with addition of ROCK inhibitor. After 11 days of dual SMAD inhibition, the cells were considered to be at NSC passage 1. The protocol was identical for both human and chimpanzee backgrounds, and three separate sets of differentiations were conducted and used as biological replicates for the CHi-C assay.

### Neuron differentiation from NSCs

The protocol for differentiating iPSCs to anterior forebrain-like excitatory neurons was based on the protocol for the STEMDiff Forebrain Neuron Differentiation and Maturation kits (catalog #s 08600 and 08605). Briefly, at day 11 post neural induction, iPSC-derived hNSCs were detached from hESC-qualified Geltrex using Accutase (STEMCELL catalog # 07920) and transferred to plates coated with Poly-L-ornithine (PLO)/laminin (Sigma Aldrich #P4957, #L2020) at low seeding density. Cells were then kept in Forebrain Differentiation Medium (STEMCELL Tech, catalog # 08601 + catalog # 08602) for 6 days for patterning into neuronal precursors and then placed in BrainPhys Neuronal Medium + Maturation supplement (STEMCELL Tech, catalog # 05797 + catalog # 08606) for up to 43 days. Neurons so produced were then detached from the plate for CHi-C and Cut and Run/RNA-Seq assays using a mix of Accutase and Papain (Worthington Co, catalog #LS003126). Two independent differentiations were conducted and used as biological replicates for the CHi-C assay.

### HiC library processing

At day 11 P1 NSCs (or day 60 neurons) were washed with DPBS (Thermofisher #14190144) and dissociated with Accutase for 5-10 minutes to obtain a single cell suspension of 20 million cells per replicate. Cells were fixed by adding 37% formaldehyde (Sigma-Aldrich #47608) to a final concentration of 2% v/v in a rotating nutator at room temperature as described previously^29^. Crosslinking was quenched using ice-cold 2M glycine (Sigma #G7126) solution to a final concentration of 250 mM. Cells were then resuspended in cold lysis buffer made by mixing 10 mM Tris-HCl pH 8 (Life Tech. # 15568-025), 0.2% v/v Igepal CA-630 (Sigma #I8896), 10 mM NaCl (Life Tech. # 24740-011), and one tablet protease inhibitor cocktail (Roche # 1187358001) and nuclear pellets were obtained by centrifugation (760xg at 4°C for 5 min). Pellets were snap frozen in liquid nitrogen and stored at -80°C. HiC libraries were prepared based on a prior Capture HiC protocol^29^. Briefly, frozen nuclei pellets were thawed on ice, digested using 50U/ul DpnII (NEB # R0543M) at 37°C overnight, followed by labeling with biotin-14 dATP (Thermofisher # 19524016) for 1 hr and ligation with T4 DNA ligase (Thermofisher # 15224025) at 16°C for 5hr. After reversing crosslinks with Proteinase K (Thermofisher # 4333793) and DNA extraction via ethanol precipitation, ligated products (50µg) were sheared by sonication (Qsonica, time=13 min, intensity=15, on-off interval=10sec) to an average fragment size of 200 bp. The ligated and sheared HiC libraries were size selected using Ampure XP beads (Beckman Coulter #A63881) and ligated fragments were pulled down using Streptavidin Dynabeads (Invitrogen #65001). This was followed by end repair and removal of biotin from non-ligated DNA ends using a mix of T4 DNA Polymerase (NeB # M0203), T4 PNK (NEB # M0201) and Klenow DNA Polymerase (NEB # M0210). After dATP tailing with Klenow fragment (3’ → 5’ exo-; NEB # M0212), we performed ligation with Illumina Dual Index UMI adaptors (NEB # E7395) and library amplification using Phusion HF PCR kit (NEB #E0553L).

### Design of CHi-C probes

We designed 120-mer DNA probes targeting DpnII fragments within a 1000bp window centered around 1590 HARs and MPRA fragments within HGEs that were called as differentially active (fragments from 466 HGEs in total were included)^5,6,8,16^. As the DpnII fragment coordinates were different in human (GRCh38) and chimpanzee (panTro6), we generated different probe sets for each genome. Each DpnII fragment initially contained 2 probes at either end, and then filtered for repeat content (BLAT density score < 40), GC content (< 60%) and duplicate sequences (<=2 duplicates) using the program Capsequm^107,108^. Final probe sequences were generated on these filtered fragments using the design tool at https://capsequm.molbiol.ox.ac.uk/cgi-bin/CapSequm.cgi and pooled custom biotinylated oligonucleotide probes for each species were ordered from Twist Biosciences Inc. The pipeline for the probe design is provided in Fig. S21.

### Capture HiC library preparation and sequencing

Target enrichment from the HiC library for CHi-C library preparation was done using the Twist Biosciences Target Enrichment Protocol with the custom biotinylated probes. Briefly, we used our indexed pooled HiC library and hybridized the probes according to the manufacturer’s instructions (Twist Fast Hybridization Kit #104180). We then bound the hybridized probes to streptavidin beads (Twist # 100983) and performed post-capture PCR amplification and DNA purification (Twist Wash Kit #104180). Then the CHi-C library was sequenced (paired-end 150bp, Illumina NovaSeq6000) for a total of 500 million reads per replicate, leading to about 1.5 billion sequenced reads each for hNSCs and cNSCs respectively, and about 1 billion sequenced reads for human neurons (Table S1).

### Capture HiC data analysis

Ligated reads were truncated, mapped and filtered to DpnII digested reference genomes (generated from the GRCh38 and panTro6 reference assemblies for human and chimpanzee respectively) using the HiCUP pipeline^48^ (truncating RE site flag = ^GATC,DpnII; mapping flags for Bowtie2: --very-sensitive, --no_unal, --reorder; ditag pairing criteria: MAPQ score >= 30, XS – AS tag ID score >= 10; filtering ditag range = 50 to 800bp). Valid ditags were deduplicated using the adapter UMIs through the umitools package^109^. Each replicate library was then evaluated for mapping %, valid ditag %, deduplication rate and cis-trans ratios as summarized in Table S1. All DpnII fragments containing HARs and HGEs were then merged into a single bait file and significant interactions were called using CHiCAGO^49^ after combining 3 replicates from the HiCUP pipeline. Briefly, we weighted the 4-parameter Delaporte model in CHiCAGO using parameters: weight(alpha) = 24.5, weight(beta) = -2.16, weight(gamma) = -21.2 and weight(delta) = -9.2, and this was used to compute weighted *P-*values (or *Q* values) for every interaction^47,110^. Non-negative scores were then assigned to each interaction on a log scale to assess significance (dependent on - log(*Q*_interaction_)). We then set a threshold for the CHiCAGO scores as >4.5 for human and >4.4 for chimpanzee, a slightly more relaxed threshold for enhancer Capture HiC than the usual 5 used for promoter CHi-C, but one which provided a significant interaction number of order 10^4^, which is in line with recommendations for CHi-C datasets^110^. The rest of the design settings for fitting the model to the data were the same as previously recommended for 4bp cutters^47^. We removed the minority of significant interactions that were *trans* in nature, i.e. the bait and interacting fragments were on different chromosomes. This gave us a set of significant *cis* interactions involving human and chimpanzee elements.

### Comparison of HAR and HGE interaction sets between human and chimpanzee

We compared the set of human interactions with the set of chimpanzee interactions for each HAR or HGE and categorized them by the proportion of common interactions. An interaction in human or chimpanzee for a given element was considered to be conserved if it fell within 10kb of the orthologous interaction in both genomes. We defined 5 categories of elements – those with no common interactions (0% conservation), along with those within 0-25%,>25-50%,>50-75% and >75-100%. To illustrate variation within each quantile, we matched the interaction coordinates of each element to obtain a distribution of conservation values in the interaction sets for the elements and plotted them as violin plots. We also defined the top decile of HARs with most conserved interactions (>90% conservation) as highly conserved HARs, and the bottom decile (0-10% conservation) as species-specific HARs.

### Annotation of CHi-C gene targets

For the classification of gene targets for the significant interactions we used the human GENCODEv43 annotation^111^. As the chimpanzee gene annotations in the panTro6 assembly were much sparser and less defined, we converted the coordinates of the chimpanzee interactions to human before assigning gene targets. Conversion was performed using the UCSC liftOver package with -minMatch =0.95. For assigning genes to HARs we used two criteria shown in Fig. S22. We first intersected the interaction coordinates from both species with the coordinates of gene annotations obtained from the GENCODEv43 GTF file, thereby yielding all genes whose gene bodies or promoters were linked to HARs, HGEs or their orthologs. We then identified HAR, HGEs and their orthologs that targeted regions 5kb upstream or downstream of annotated promoters and considered these as gene targets. The final gene target sets were derived from the union of both approaches. GO enrichment analysis of gene target sets were performed using the EnrichR package^112^ (using all protein-coding genes as the background) and an FDR *Q*-value cutoff of 0.05.

For the classification of element interactions with active or repressed promoter regions shown in Fig. 3F, we used the histone modification profiles we generated via CUT&RUN (described below). We defined active genes based on H3K27ac occupancy at their promoters, while repressed genes were defined by H3K27me3 occupancy at their promoters. Putative enhancers were called as H3K27ac or H3K27me3 marked regions outside of promoters that were located in introns or intergenic regions. Interactions potentially mediated by cohesin/CTCF involving HARs and HGEs were identified based on the occupancy of CTCF and RAD21 within the interacting region intervals (+/- 500bp). For comparing differences of chromatin marking (or ENCODE cCRE presence) at conserved versus species-specific interactions, we used a 2x2 contingency table (interaction type: conserved, species-specific as columns, chromatin marking categories: any mark, no mark as rows) and performed a Fisher’s exact test, using a *P-*value cutoff of 0.05 to call significance.

For the breakdown of the CHi-C interactions across the genome shown in Fig. 3E, promoters (+/- 2kb of TSS), intronic and intergenic regions were characterized based on GENCODEv43 annotation. Non-exonic constrained elements defined by phastCons (called on the UCSC Multiz 100-way vertebrate alignment) were intersected with the interaction coordinates for both species to measure the constraint of regions interacting with these elements. HAR-to-HAR and HGE-to-HGE interactions were defined reciprocally, i.e. each HAR or HGE in the pair was required to interact with the other HAR or HGE.

### Analysis of HAR and HGE contact regions for interaction profiles

The interaction profiles of each HAR and HGE were visualized by plotting the significance of its interactions against the relative distance of the interacting region from the element (e.g., Fig.2B). In many cases due to the high-resolution nature of the CHi-C assay we identified regions with a high density of significant interactions forming peak-like distributions, so we plotted a density distribution to find regions of high interaction density.

Contact regions for the density scatterplots were defined by using the localized peak of the density distribution. A kernel density distribution was used to define a peak at every cluster in the neighborhood of each HAR or HGE. A custom script was used to define the contact regions as a neighborhood on either side of the local maxima determined by the full width half maximum (FWHM) of local peaks for the entire interaction profile of the HAR or HGE. This allowed us to identify the contact regions shown in Fig. 2 and Fig. S2.

### Comparison of other published HiC datasets with results of this study

The number of HARs sampled and number of gene targets for three of the four published studies were listed in Table 1 as reported in each study^18,26,28^. As the fourth study (Keough *et al*.) did not provide a comprehensive list of gene targets^7^, we used the HiC interaction coordinates provided by the study to identify interactions for zooHARs from that study and HARs analyzed here, and then used the criteria described above with the GENCODEv43 annotation to identify gene targets.

We used a permutation approach as described in Keough *et al.* to determine whether HARs, HGEs, or VISTA enhancers were significantly enriched within TADs containing hsSVs (*P_adj_* < 0.05). Briefly, we determined whether zooHARs, VISTA enhancers, HARs, and subsets of HARs with highly conserved or species-specific interaction profiles (top and bottom deciles as described in the Results) were enriched in TADs called in the germinal zone and cortical plate of the fetal human brain that also had at least one hsSV^25,53^. Using the approach described in Keough *et al.*, we then compared the observed number of elements from each category that were within hsSV-containing TADs to a background distribution constructed by randomly selecting an equal number of phastCons elements 20,000 times and determining how many fell with hsSV-containing TADs for each trial. Significance of enrichment for each category was then evaluated based on the proportion of permutation trials that showed a greater number of phastCons elements assigned to hsSV-containing TADs than the observed number for each category. We also used this permutation approach to test the enrichment of HGEs in TADs with hsSVs compared to a background distribution constructed by drawing from any genomic element showing H3K27ac or H3K4me2 activity across fetal human brain samples described in Reilly *et al*.^22^ (n = 20,000 trials).

### CUT&RUN sample processing

CUT&RUN was performed on human and chimpanzee NSCs (P1 each) as well as human iPSC-derived neurons for 5 separate chromatin marks (H3K27ac, H3K27me3, CTCF, RAD21, POLII) and 2 replicates each in one batch using Epicypher CUTANA CUT&RUN kit (SKU:14-1048) according to manufacturer’s instructions. Libraries were multiplexed and sequenced with paired end 150bp reads on an Illumina NovaSeq6000 sequencer.

### CUT&RUN data analysis

Reads were trimmed for adapters using Trimmomatic^113^ and aligned to either the GRCh38 (human) or panTro6 (chimpanzee) genomes using Bowtie2 (flags: --very-sensitive, -X 2000)^114^. Aligned reads were filtered based on quality and PCR duplicates were marked using the MarkDuplicates function in Picard (courtesy the Broad Institute bioinformatic toolkit)^115^. Reads with the following flags – “reads unmapped”, “not primary alignment reads” and “failing platform duplicates” were removed and the remaining properly mapped reads were used for peak calling using the stringent setting of the program SEACR^116^, which was used to call enriched regions in target data by selecting the top 0.5-1% of regions by AUC based on the assayed mark. Consensus peaks for each mark were called by intersecting reproducible peaks from both replicates. The signal in the BigWig files for visualization was obtained through the deeptools bamCoverage package^117^, normalized to read depth (--normalizeUsing CPM --binSize 10 --extendReads 300 -- centerReads). Active and repressed interactions from each cell type were classified in a bidirectional manner, where both the HAR or HGE and its gene target had to be located inside a 500bp window around the respective consensus peak (H3K27ac for activating interactions, H3K27me3 for repressive interactions). We classified switches from repressed interactions at the NSC stage to active interactions at the neuron stage if the HAR/HGE-gene target pair was located near a H3K27me3 consensus peak in hNSCs (flag for bedtools slop: -b 500bp) and the corresponding interaction was near a H3K27ac consensus peak in neurons (flag for bedtools slop: -b 500bp, bidirectional).

### RNA-Seq data generation and analysis

Cells were collected from four samples of human and chimpanzee NSCs at early passage (P2) in one batch, and from three samples of human iPSC-derived neurons. RNA isolation was done using the Qiagen RNeasy Plus kit and library preparation and sequencing was performed by the Yale Center for Genome Analysis. For the hNSC versus cNSC comparison, paired-end reads (100bp) were mapped to a modified version of the XSAnno whole transcriptome annotation^118,119^, which integrates the orthologous regions of human and chimpanzee transcripts so we can do inter-species comparisons. This consensus annotation was built on the GENCODEv43 annotation for the GRCh38 and panTro6 genomes. Briefly, we subsetted out genes with exons that reciprocally lifted over between both species (>98% sequence similarity between GRCh38 and panTro6 genomes), then filtered out those with ambiguous annotations using BLAT [blat parameters: % ID between species = 95%, % len between species = 95%, % ID within species (paralogs) = 97%, % len within species = 97%] and finally simulated reads using simNGS to filter out exons that were differentially expressed in the simulation using DESeq2^120^. Modifications to the workflow and additional scripts we used have been deposited on GitHub. Mapping was performed using the STAR aligner (flags: --outSAMunMapped Within, --twopassMode Basic, -- outFilterMultimapNmax 1, --quantMode TranscriptomeSAM)^121^. Using the package RSEM^122^ we generated a counts matrix for each species’ replicate and then used DESeq2 for differential gene expression analysis. Genes with FDR-adjusted *P*-value < 0.01 and |log2FC| > 1 were considered differentially expressed between human and chimpanzee (Table S2). For the hNSC-neuron comparison, reads were mapped to native GRCh38 coordinates and TPM counts (RefSeq annotation) in each cell type were compared for specific gene targets shown in Fig. 5.

### Human and chimpanzee multiplexed single-cell RNA-seq

Human and chimpanzee iPSCs underwent dual SMADi neural induction for 10 days as described above, then were collected separately in 1x DBPS with 0.04% BSA (Thermo Scientific #B14). Each sample was labeled using 3’ CellPlex Kit Set A (10X Genomics cat #100261), then pooled. Approximately 10,000 cells were profiled for scRNA-seq across 2 GEM lanes on a 10X Chromium Controller using Chromium Single Cell 3’ v3.1 Reagents with Feature Barcoding (PN-1000268). Library preparation and sequencing was performed by the Yale Center for Genome Analysis. Reads were aligned to GRChg38 using the *multi* function of CellRanger 6.0.1^123^. For quality control, cells with either nFeature_RNA (number of genes) or nCount_RNA (number of molecules) values 2 standard deviations away from the mean were removed, alongside cells with mitochondrial reads greater than 10% of total reads. Counts were normalized using Seurat (v4.3.0)^124^, via the NormalizeData function with default parameters. Human and chimpanzee NSC count matrices were then integrated using the SelectIntegrationFeautres, FindIntegrationAnchors, and IntegrateData functions with default parameters. Dimensionality reduction was performed using RunPCA (params: npcs = 30) and RunUMAP on the integrated data, and Louvain clustering was performed using FindNeighbors (params: dims = 1:15) and FindClusters (params: resolution = 0.3). Cluster identities were assigned by identifying cell-type markers using FindAllMarkers (params: min.pct = 025). Cell type clusters identified are: ECM-vRG – extracellular matrix/ventricular radial glia like (marker genes *TAGLN, ANXA1, ANXA2, CAV1* and *ACTN1*), oRG like – outer radial glia like (marker genes *PTPRZ1, FAM107A* and *HOPX*), Cycling RG – dividing radial glia/progenitor like (marker genes *TOP2A, SNHG19* and *MKI67*), IP – intermediate progenitors (marker genes *EOMES, ELAVL4* and *NEUROG1*) and NC – neural crest (marker genes *BMP7, MSX1, PAX3* and *SNAI2*).

### Differential marking (DM) analysis

The SEACR peaks for each chromatin mark (H3K27ac and H3K27me3) and each species (human and chimpanzee) were combined using the following procedure: reproducible peaks from both replicates were intersected to identify the largest interval that encompassed the peak regions, which we referred to as consensus regions. Subsequently, the consensus regions for a particular mark were merged across both species and reciprocally lifted over between both genomes with liftOver -minMatch threshold of 0.9. This step aimed to obtain consensus regions that covered the entire set of orthologous enriched peaks across both species for that specific mark. The resulting common consensus region BED file was then converted to a simple annotation text format (SAF). To determine the raw counts of paired-end reads present within these consensus regions for both replicates of each species, we utilized the featureCounts package from the Subread module^125^ with the SAF file for the -a flag. Size factors were estimated using the --normalizeUsing CPM flag in the deeptools package, and differential enrichment for that mark between the two species was computed using DESeq2. A consensus region was considered differentially enriched (DM) if the adjusted *P*-value (FDR) assigned by the model was < 0.01. If the |log2FC| was > 1, the consensus region was termed differentially marked (DM), and are provided in Table S2.

Finally, the coordinates of all differentially marked peaks were intersected with the HAR/HGE coordinates. This intersection provided a list of HARs/HGEs and their orthologs that exhibited a human or chimpanzee-specific chromatin mark. The presence and absence of species-specific H3K27ac and H3K27me3 peaks at these elements were then associated with the ratio of TPM expression values of the genes from human and chimpanzee NSCs obtained from the XSAnno mapping. To find the curve of best fit for H3K27ac deposition at the HAR vs gene target expression, we used a residual-square error (RSE) estimator function, which computed the polynomial regression (for all polynomials of order < 5) with the minimum cumulative MSE for the points in the point cloud. Using this metric, we estimated that a third degree polynomial fit the point cloud the best, and used the stat_smooth() function in ggplot to fit a model y∼O(x^3^). Pearson’s product-moment correlation was used to compute the correlation between the y and O(x^3^) variables. A similar estimation for the H3K27me3 case gave a curve of best fit as y∼O(x^2^), and we computed the Pearson correlation between the y and O(x^2^) variables.

### Over-representation of DM HARs/HGEs with DEG targets

We used a permutation approach to ask whether the DM HARs/HGEs were enriched for at least 1 DEG target if they are significant at 5% *P*-value level for the following test: are DM HARs/HGEs enriched for DE genes compared to an empirically constructed background distribution? Briefly, the background set of genes was the set of HARs/HGEs with H3K27ac signal in both species and containing a gene target, and randomly sampled element sets (of the same size as the DM HAR/HGE set) from this background were intersected with the set of all HARs/HGEs with a DEG target for a total of 20,000 trials to generate the empirical background distribution. Then the *P*-value was computed as the proportion of trials greater than the observed value between the DM HAR/HGE target set with at least 1 DEG target.

### Enrichment of all HAR and HGE gene target sets within differential gene sets

Gene lists were taken from the set of human-specific differentially expressed genes (hDEGs) for all cortical cell types provided by Jorstad et al^32^ and the set of human-specific CRE linked genes (hDAGs) provided by Caglayan et al^33^ respectively. We used a permutation approach to ask whether the gene target set of HARs/HGEs are enriched near hDEGs or hDAGs if they are significant at 5% *P*-value level to determine if HAR targets were enriched for genes called as hDEGs or hDAGs compared to an empirically constructed background distribution. Briefly, the background set of genes was the set of genes in the annotations used by each paper, and randomly sampled gene sets (of the same size as the HAR gene target set) were intersected with the hDEGs and hDAGs for a total of 20,000 trials to generate the background distribution. Then the *P*-value was computed as the proportion of trials greater than the observed intersection between the HAR target set and the corresponding human-specific gene set and was corrected using the Bonferroni procedure based on the number of tests run. An identical permutation approach was taken to compute enrichment of HAR and HGE gene targets within DEG sets identified between human and Rhesus macaque prenatal brains across at least one of multiple brain regions sampled, as well as within individual regions such as the pre-frontal cortex^34^. We also used this permutation approach to identify WGCNA modules with an enrichment of HAR/HGE gene targets, using FDR correction for multiple tests and identified 15 enriched modules. GO enrichment analysis for each enriched module was performed using the EnrichR package^112^ (using all genes used to compute WGCNA modules as the background) and an FDR *Q*-value cutoff of 0.05. The Main GO Term was defined for the module if any daughter term under a collapsed heading in ReviGO^126^ (scale set to 0.4) matched one of the five broad categories set out in Fig. 3D.

### Enrichment of conserved and species-specific HAR and HGE gene targets within differentially expressed gene sets

We then performed enrichment tests in an identical fashion for the conserved and species-specific gene targets for hDEGs, hDAGs, and DEGs called between human and macaque prenatal brain regions as described above, as well as the DEGs identified in hNSCs and cNSCs in this study, and a set of DEGs identified between human and chimpanzee hippocampus intermediate progenitors generated from the same iPSC lines used in this study^68^. Briefly, for the conserved or species-specific gene target sets, we generated two background distributions comprised of the number of intersections of 20,000 randomly sampled gene sets of the same size as either the conserved or species-specific gene target set with each DEG or DAG set. Relative enrichment of the conserved set compared to the species-specific set for each test was computed as Enr = O_1_/O_2_, where O_1_ is the ratio of the observed intersection of the conserved gene target set and the differentially expressed gene set and the mean of the background distribution generated for the conserved set, and where O_2_ is the ratio of the observed intersection of the species-specific gene target set and the differentially expressed gene set and the mean of the background distribution generated for the species-specific set. *P* values were computed as the number of values of each background distribution greater than the observed value for the corresponding gene target set and corrected using the Bonferroni procedure based on the number of tests run.

### CRISPRa and CRISPRi validation experiments

To validate the predictions from the integrated CHi-C and Chromatin maps we chose one active and one repressed loop involving a HAR and its contact genes and used CRISPRi and CRISPRa to repress or activate the HAR, respectively, and then determined if there was any impact on target gene expression. CRISPR guides for HARs were designed along an orthologous region in both human and chimpanzee with low off-target binding scores (determined by the program FlashFry^127^; thresholds used: hsu_value >= 65.0, jost_ot <= 0.4, jost_spec >= 0.5, gc_flag = ‘NONE’, polyT_flag = ‘NONE’, mismatch_0 = 1 and mismatch_1 = 0 and mismatch_2 = 0). We used CRISPRa and CRISPRi to respectively active or repress *HACNS52* (part of an H3K27ac-marked active loop with *ANXA2* and *ICE2*) in human and chimpanzee NSCs, using two independent guides. The 20bp guide target sequences were 5’AGTATTTGTGTTCTCCGGAA 3’ and 5’ TCTAGTTATTTGGACCCATA 3’. For each CRISPR experiment, the guides were cloned into the respective CRISPRa and CRISPRi vectors from Origene Tech (CRISPRa vector has a tGFP reporter, dCas9-VP64 and a gRNA cloning site, cat # GE100074; CRISPRi vector has a tGFP reporter, dCas9-KRAB-MeCP2 and a gRNA cloning site, cat # GE100085) according to manufacturer’s instructions, and transfection was carried out using Turbofectin 8.0 reagent (cat # TF81001, Origene Tech). Two replicates were conducted for each CRISPRa and CRISPRi experiment. Transfection efficiency was measured using GFP expression 48 hours post transfection in the cells on a CellDrop fluorescence cell counter (DeNovix, model FL); efficiency in all cases was >90%. Scrambled guides for both CRISPRa (cat # GE100077) and CRISPRi (cat # GE100086) from Origene Tech were included as controls. The effect of HAR perturbation was measured by performing qPCR on the perturbed vs control human and chimpanzee NSCs for gene target expression (relative to the reference gene *TBP*). Fold change was computed using ΔΔCT for each perturbation, and statistical significance was determined using a 2-way ANOVA test (α = 0.05) followed by a Tukey HSD Test (*P < 0.1*).

### Characterizing cell type-specific expression across progenitor and neuronal subtypes from eight regions of the developing human brain

Single cell RNA sequencing (scRNA-seq) datasets collected from multiple regions of the developing human brain were published alongside two brain atlas publications^35,36^. We acquired processed counts matrices from seven forebrain regions (cortex, hippocampus, MGE, LGE, CGE, thalamus, hypothalamus) and the cerebellum at time points ranging from CS14 to GW25. These datasets were loaded into the Seurat R package (v4.3.0)^124^ and each dataset was filtered independently to keep only cells with a minimum of 750 represented features and features with expression in at least 50 cells. Datasets with fewer than 100 cells after filtering were considered low quality and discarded. Additionally, cells with read or feature counts greater than two standard deviations from the mean and cells with greater than 10% mitochondrial read percentage were considered doublets, empty, or dying cells and were filtered out.

Resulting high-quality cells in each dataset were then merged into a single Seurat object per region for a total of eight Seurat objects. The subsequent pipeline was performed for each of these Seurat objects respectively. The counts matrix was normalized (NormalizeData), subset to the top 2000 highly variable features (FindVariableFeatures), and then scaled (ScaleData). The merged datasets were then integrated with FastMNN^128^ (RunFastMNN in SeuratWrappers v0.3.0, params: k = 30, d = 50) to remove batch-related technical variation. We then calculated the 30 nearest neighbors (FindNearestNeighbors, params: k = 30, reduction = ‘mnn’) for each cell using the 50 integration features calculated from FastMNN. These integration features were then used to cluster transcriptomically-similar cells with the Louvain neighborhood aggregation algorithm in Seurat (FindClusters, params: resolution = 0.85). Marker genes for each cluster were calculated in Seurat (FindAllMarkers, params: min.pct = 0.25, only.pos = TRUE). Clusters were first partitioned into progenitors (*SOX2, MKI67, MCM3*) and neurons (*STMN2, DCX*). Granular cell type assignments were made for clusters with expression profiles that reflected known subtypes (e.g. *TBR1*+ cortical deep layer neurons) or, in cases where no clear cell type markers emerged, clusters were assigned generic regional classifications. Progenitor populations born outside of the cortex but expressing *EOMES* or *HOPX* were classified as NSC2 or NSC3 respectively. Additionally, the cell cycle phase of each progenitor cluster was assigned using the expression of *TOP2A* and *MKI67* marking G2/M cells.

Neural stem cell or neuronal clusters were then extracted and merged independently. The following pipeline was then performed for the neural stem cell and neuron populations independently. Merged datasets were integrated using the scArches Python package (v0.5.7)^129^, resulting in integration features that were used to calculate UMAP coordinates with the scanpy Python package (v1.9.3)^130^ for downstream visualization. Each cell type was downsampled to a maximum of 2000 cells and the normalized expression values were then scaled in Seurat (ScaleData). We extracted the average expression value of each HAR contact across all progenitor or neuronal cell types using Seurat (AverageExpression), resulting in cell-type-by-gene matrices of average scaled gene expression values. We clustered these expression values for each gene using the k-means algorithm in the stats R package (kmeans, params: centers = 15). The average expression values for each cluster were visualized separately using ComplexHeatmap^131^ and labeled with their respective cell type-specific expression profiles. Clusters with relevant expression patterns were isolated and visualized with ComplexHeatmap. For progenitor populations, the cell cycle phase for each cell type was annotated alongside the expression profiles. Expression profiles of candidate genes for each cluster were visualized on the UMAP embedding using density plots (plot_density from Nebulosa v1.2.0 R package). Additionally, average scaled expression profiles were extracted for human-biased DEG HAR contacts in neural stem cells and reclustered using kmeans (params: centers = 3), revealing a cortical oRG-biased cluster visualized using ComplexHeatmap as described above.

