## Supplemental Figures for "Resolving the three-dimensional interactome of Human Accelerated Regions during human and chimpanzee neurodevelopment"

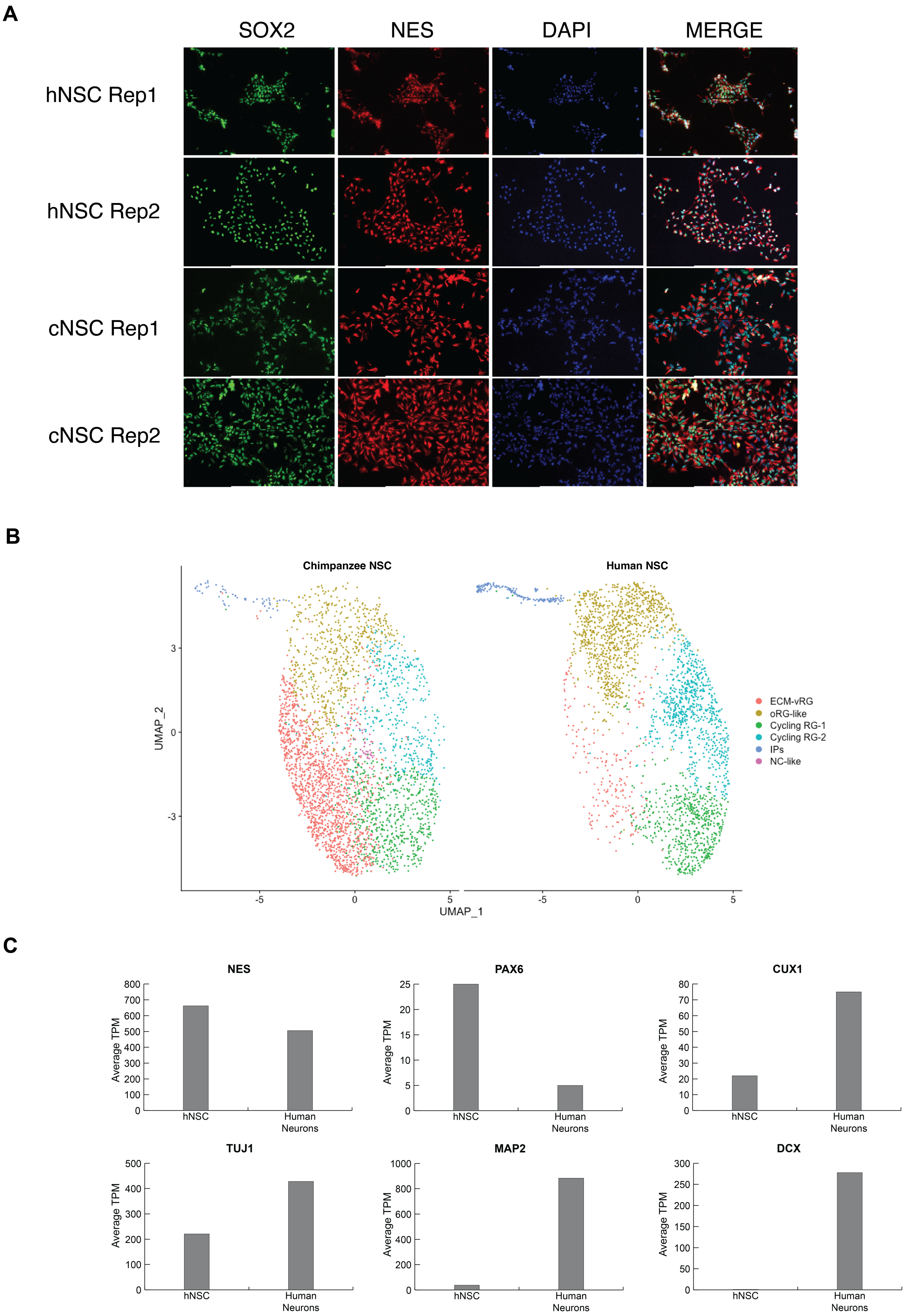

A
SOX2
NES
DAPI
MERGE
hNSC Rep1
hNSC Rep2
cNSC Rep1
cNSC Rep2
B
Chimpanzee NSC
Human NSC
UMAP_2
UMAP_1
ECM-vRG
oRG-like
Cycling RG-1
Cycling RG-2
IPs
NC-like
C
NES
PAX6
CUX1
TUJ1
MAP2
DCX
Average TPM
hNSC
Human Neurons

**Supplementary Figure 1**. **Characterization of human and chimpanzee neural stem cells and human neurons.** **(A**) Expression of NSC markers in human and chimpanzee NSCs (hNSCs and cNSCs respectively), visualized using immunofluoresence. SOX2, green*;* NES, red; DAPI nuclear staining, blue. **(B)** Single cell RNA-sequencing of hNSC and cNSC populations (UMAP visualization). Cell types abbreviations are: ECM-vRG – extracellular matrix/ ventricular radial glia like, oRG like – outer radial glia like, Cycling RG – dividing radial glia/progenitor like, IP – intermediate progenitors and NC – neural crest. Marker genes used to identify each cell type are provided in the Methods. **(C)** Expression profiles (from RNA-Seq) of known NSC marker genes (*NES, PAX6*) and neuron marker genes (*CUX1, TUJ1, MAP2, DCX)* in hNSCs and neurons used in this study.

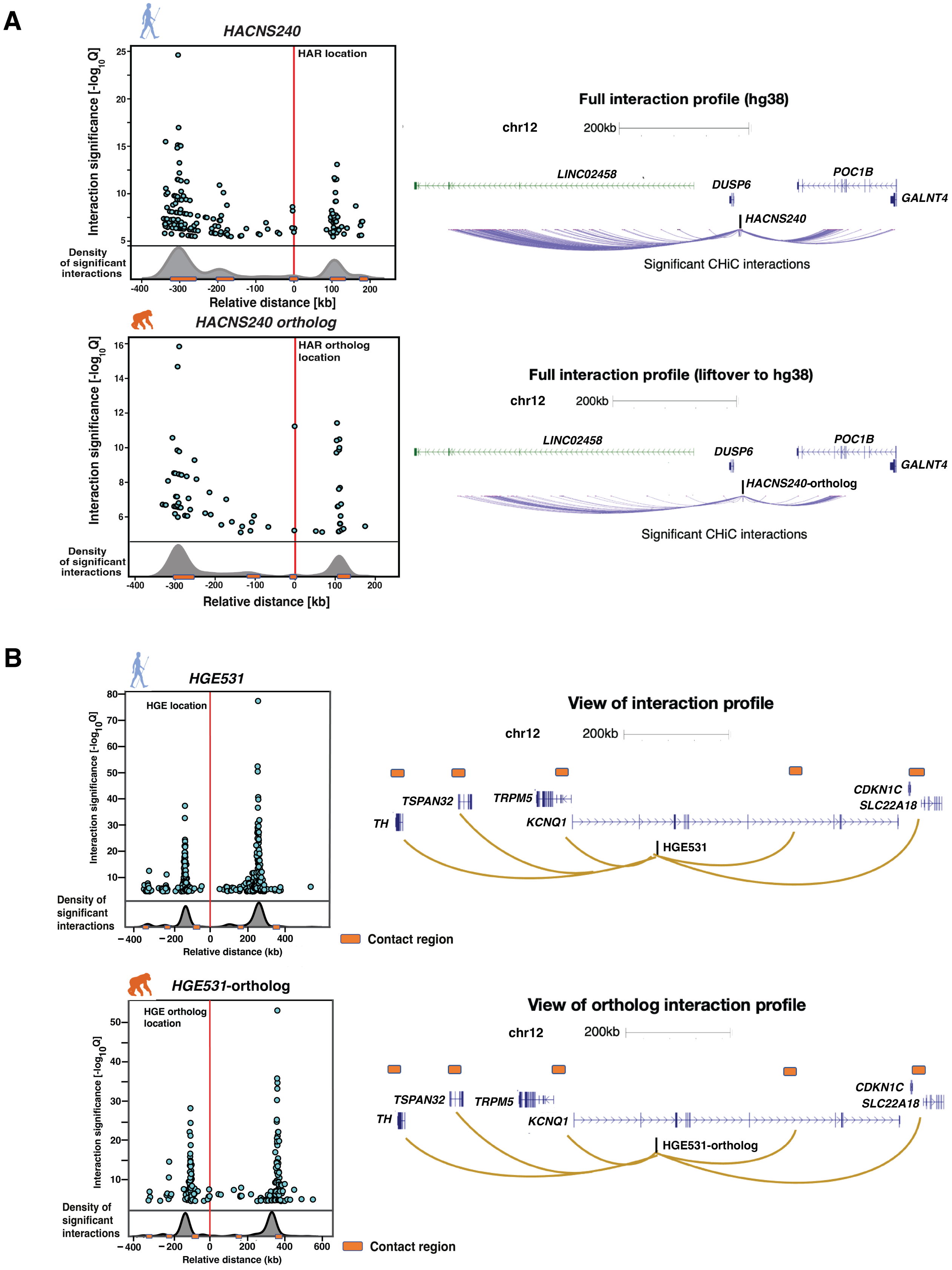
A
HACNS240
HAR location
Interaction significance [-log10 Q]
Density of significant interactions
Relative distance [kb]
Full interaction profile (hg38)
chr12
200kb
LINC02458
DUSP6
POC1B
GALNT4
HACNS240
Significant CHiC interactions
HACNS240 ortholog
HAR ortholog location
Full interaction profile (liftover to hg38)
HACNS240-ortholog
B
HGE531
HGE location
Relative distance (kb)
Contact region
View of interaction profile
TSPAN32
TRPM5
TH
KCNQ1
CDKN1C
SLC22A18
HGE531-ortholog
HGE ortholog location
View of ortholog interaction profile

**Supplementary Figure 2**. **Interaction profile examples of HARs and HGEs in hNSCs and cNSCs. (A)** *Left.* Detailed interaction profiles for *HACNS240* (*top)* and its chimpanzee ortholog (*bottom)*, as in Fig. 2B. Each point on the scatterplot represents a significant interaction, plotted by the degree of significance as shown on the y-axis. The relative distance from the HAR is given on the x-axis. A normalized density distribution of the interactions is shown below the scatterplot, and regions of high interaction density are marked by orange boxes (Methods). *Right.* Schematized UCSC Genome Browser view (in GRCh38 coordinates) of the interaction distribution for all significant interactions in both human and chimpanzee NSCs (shown as purple). Protein-coding genes are shown in blue, and lncRNA genes are shown in green. **(B)** *Left.* CHiC interaction profiles of HGE531 (*top*) and its chimpanzee ortholog (*bottom*). The plots are the same as described in **(A)**. *Right*. Schematized UCSC Genome Browser view (in GRCh38 coordinates) for each interaction profile. The location of HGE531 and its ortholog are shown, and the regions of high interaction density are shown in orange. Representative looping events are shown in gold. Protein-coding genes are shown in blue.

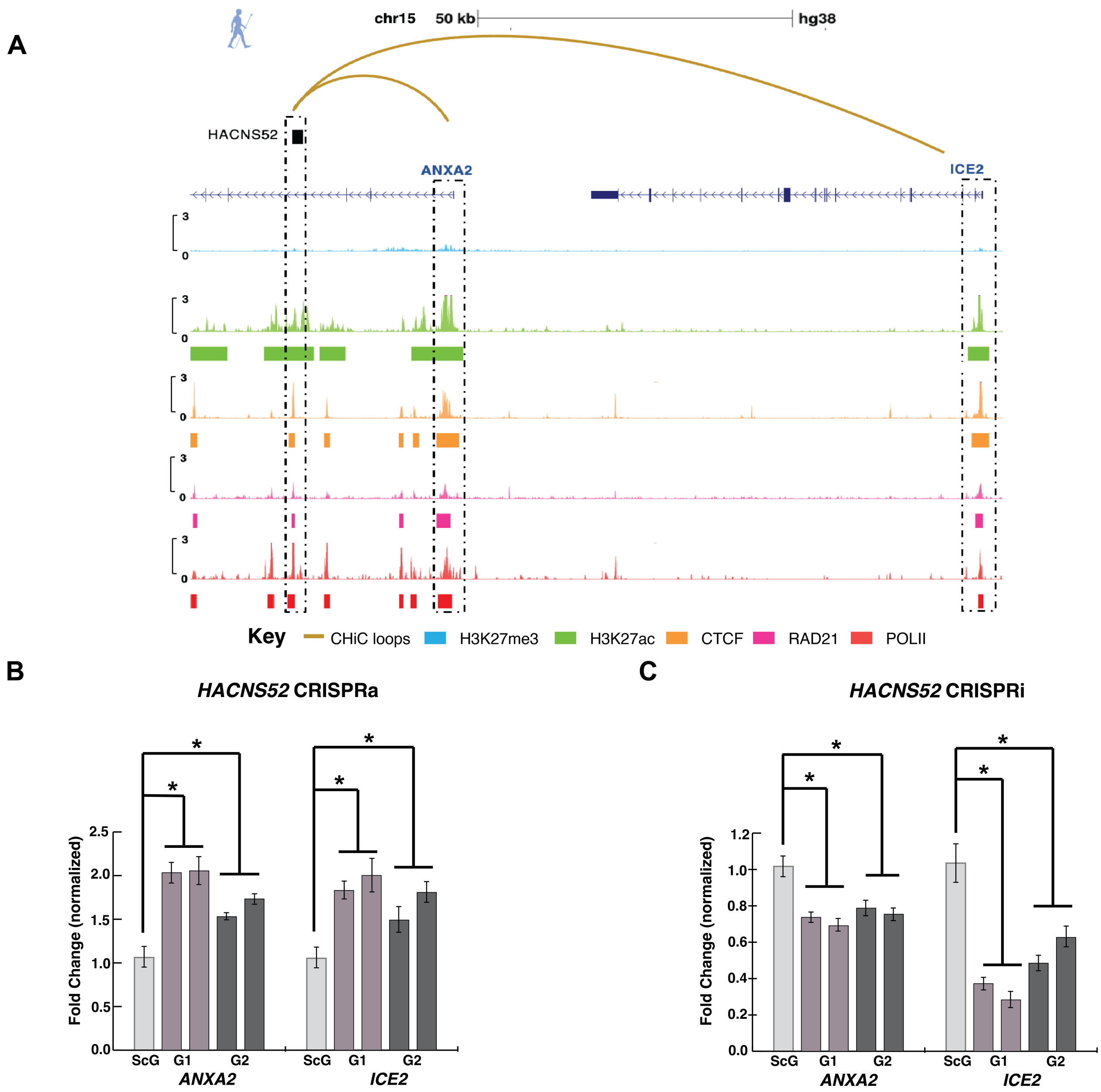


**Supplementary Figure 3. Interactions involving HARs can affect gene target expression in hNSCs. (A)** Region containing *HACNS52* (GRCh38 coordinates), which forms an active loop with its gene targets *ANXA2* and *ICE2* in hNSCs. Curved golden lines indicate representative CHiC interactions involving the HAR, and the following CUT&RUN signal tracks are shown from top to bottom: H3K27me3 in blue, H3K27ac in green, CTCF in orange, RAD21 in pink and POLII in red. The overlap between CUT&RUN signal, the HAR and its gene targets is highlighted by dashed black boxes. Data were visualized via the UCSC Genome Browser. **(B)** Effect of *HACNS52* activation (CRISPRa) on gene target expression in hNSCs using two independent guides (G1, G2). Fold-change values are measured by qRT-PCR with respect to gene expression for scrambled guide (ScG), and significance computed using 2-way ANOVA ($\alpha = 0.05$) followed by Tukey HSD test

(* indicates $P_{adj} < 0.1$ in the Tukey HSD test). **(C)** Effect of *HACNS52* inactivation (CRISPRi) on gene target expression in hNSCs using two independent guides (G1, G2). Fold-change values are measured by qRT-PCR with respect to gene expression for scrambled guide (ScG), and significance is computed in the same manner as in **(B)**.

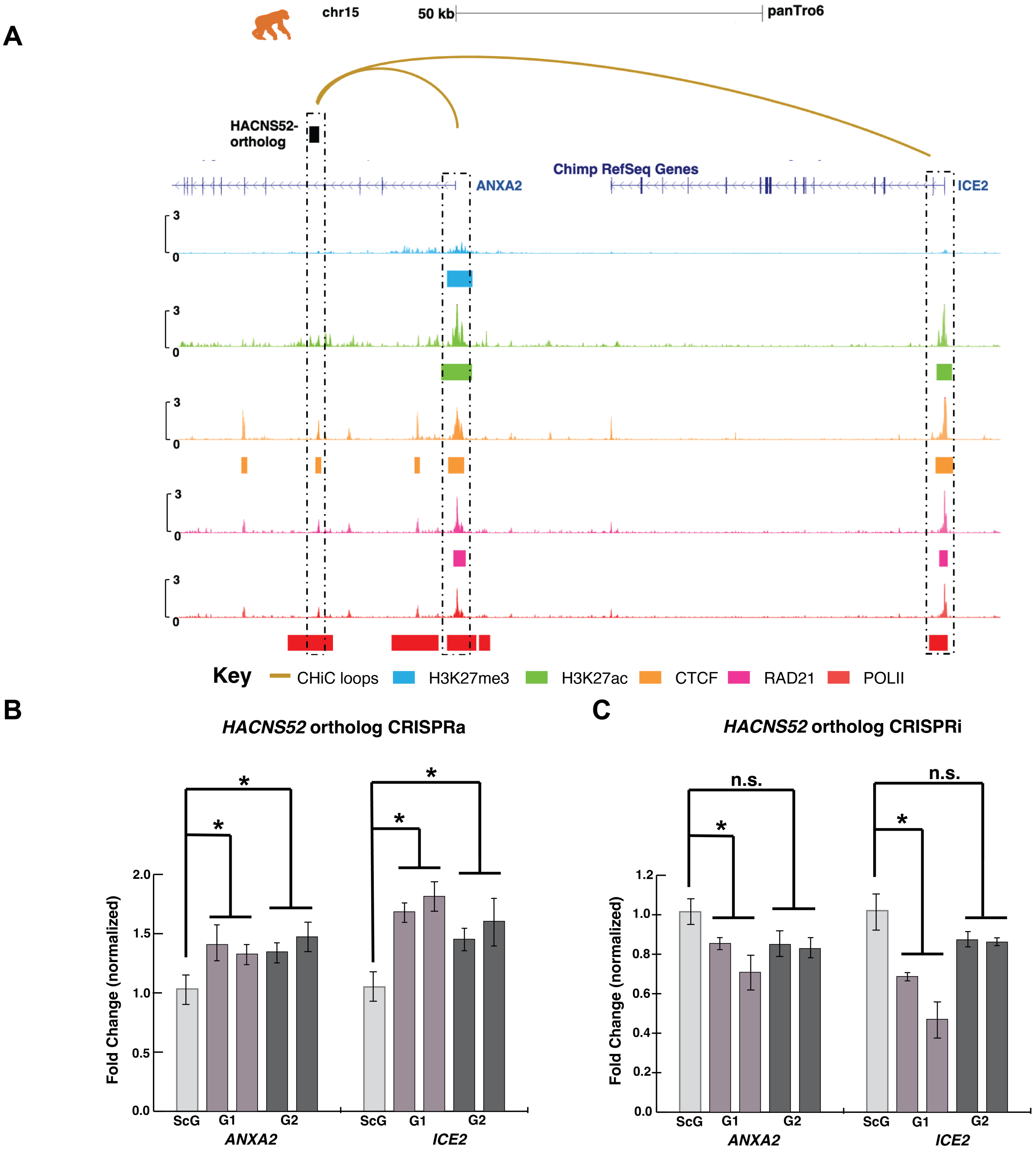


**Supplementary Figure 4. Interactions involving HAR orthologs can affect gene target expression in cNSCs. (A)** Region containing the chimpanzee ortholog of *HACNS52* (PanTro6), which forms an active loop with its gene targets *ANXA2* and *ICE2* in cNSCs. CHiC interactions and CUT&RUN signal tracks are shown in the same manner as in Fig. S3A**.** Data were visualized via the UCSC Genome Browser. **(B)** and **(C)** Effect of *HACNS52*-ortholog activation (CRISPRa) and inactivation (CRISPRi) respectively in cNSCs using two independent guides (G1, G2). Fold-change values are measured by qRT-PCR with respect to gene expression for scrambled guide (ScG), and significance is computed in the same manner as in Fig. S3B and Fig. S3C**.**

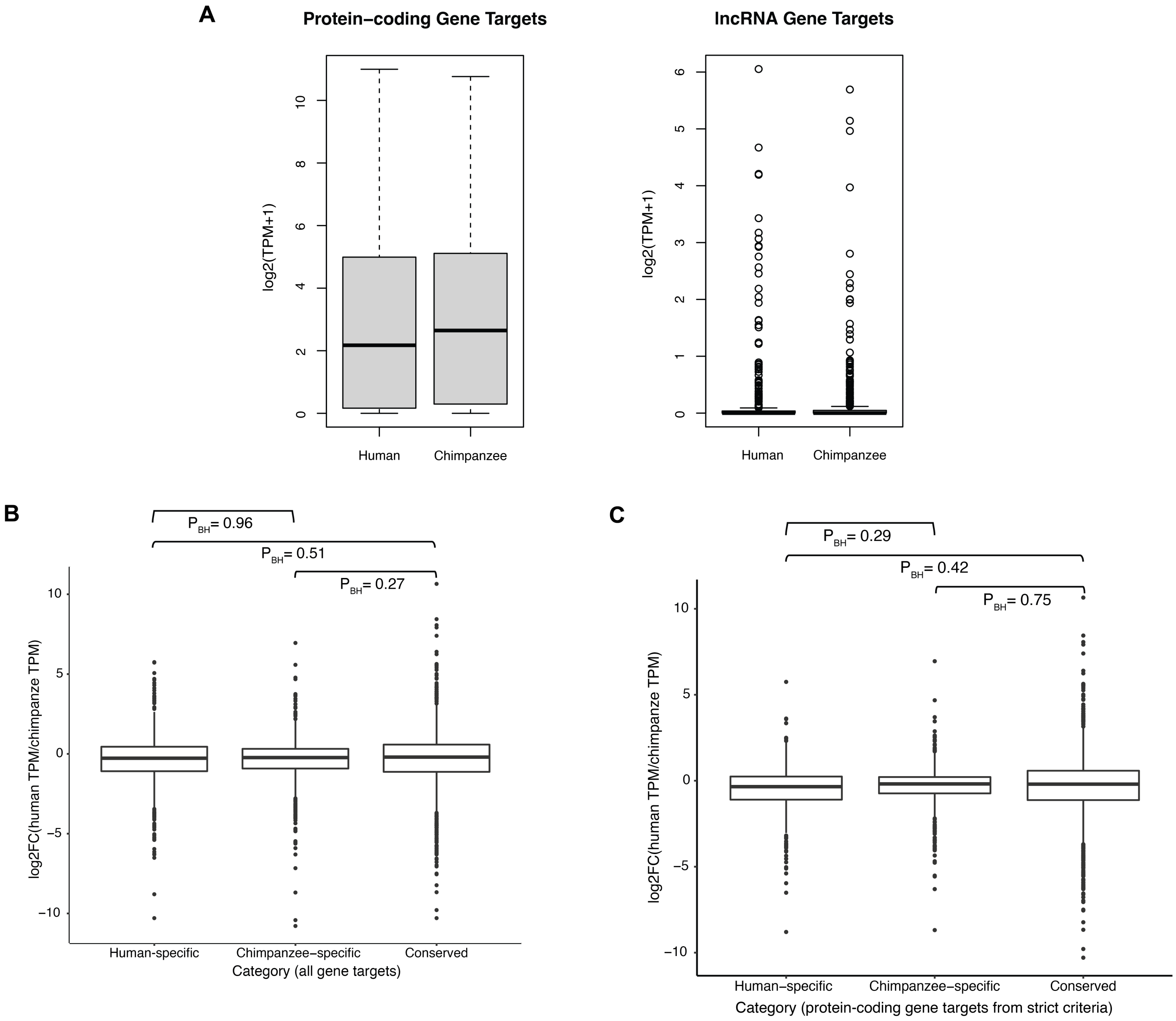


**Supplementary Figure 5. Expression profiles of different categories of HAR and HGE gene targets.** **(A)** Distribution of gene expression (log2(TPM+1)) of all protein-coding and lncRNA gene targets in hNSCs and cNSCs. **(B)** Distribution of relative gene expression of conserved set of gene targets versus human-specific and chimpanzee-specific gene targets in hNSCs and cNSCs (values provided are log2(human TPM/chimpanzee TPM). *P*-values are computed from a Wilcoxon test and corrected using the BH procedure ($P_{BH}$). **(C)** Distribution of relative gene expression of a refined list of conserved gene targets versus human-specific and chimpanzee-specific gene targets, defined using the strict criteria in Fig. S22, in hNSCs and cNSCs. Values provided are log2(human TPM/chimpanzee TPM), and $P_{BH}$ values are computed as in **(B)**.

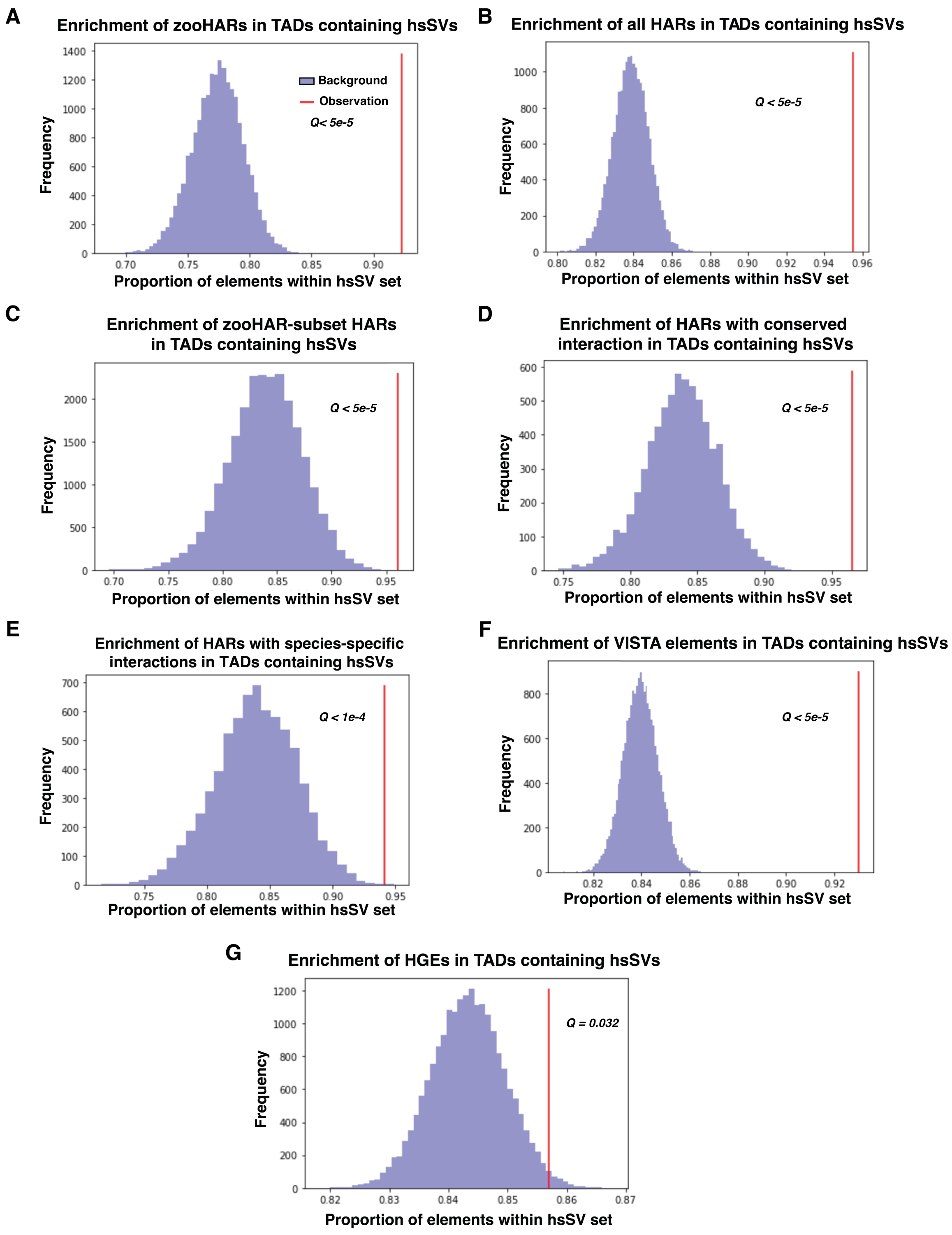
A
Enrichment of zooHARs in TADs containing hsSVs
Background
Observation
Q< 5e-5
Frequency
Proportion of elements within hsSV set
B
Enrichment of all HARs in TADs containing hsSVs
Q < 5e-5
Frequency
Proportion of elements within hsSV set
C
Enrichment of zooHAR-subset HARs in TADs containing hsSVs
Q < 5e-5
Frequency
Proportion of elements within hsSV set
D
Enrichment of HARs with conserved interaction in TADs containing hsSVs
Q < 5e-5
Frequency
Proportion of elements within hsSV set
E
Enrichment of HARs with species-specific interactions in TADs containing hsSVs
Q < 1e-4
Frequency
Proportion of elements within hsSV set
F
Enrichment of VISTA elements in TADs containing hsSVs
Q < 5e-5
Frequency
Proportion of elements within hsSV set
G
Enrichment of HGEs in TADs containing hsSVs
Q = 0.032
Frequency
Proportion of elements within hsSV set

**Supplementary Figure 6**. **Enrichment of several enhancer classes within TADs containing human-specific structural variants. (A- C)** Enrichment of zooHARs (**A**), HARs (**B**) and the subset of HARs in this study that are zooHARs (**C**), respectively, within TADs containing human-specific structural variants (hsSVs), compared to randomly chosen phastCons conserved elements. The TADs were defined in the germinal zone of the brain (containing neural progenitors) by Won *et. al* [28], and we used the hsSVs referenced in Keough *et. al* [9,45]. The enrichment *Q*-values were computed based on random sampling of the background (n = 20,000 trials; Methods). **(D, E)** Enrichment test to show enrichment of HARs with the most conserved interactions (top 10% most conserved), and HARs with the most species-specific interactions (top 10% least conserved) respectively, within TADs containing hsSVs, compared to randomly chosen PhastCons conserved elements. TADs, hsSVs and *Q*-value computation were the same as in **(A)**. **(F, G)** Enrichment of enhancer elements from the VISTA database and HGEs in TADs containing human-specific structural variants (hsSVs). VISTA elements were compared to a background of randomly chosen phastCons conserved elements, while HGEs were compared to a background of other regions showing enriched H3K27ac or H3K27me2 deposition in the fetal human brain[21]. TADs, hsSVs and *Q*-value computation were identical to **(A)**.

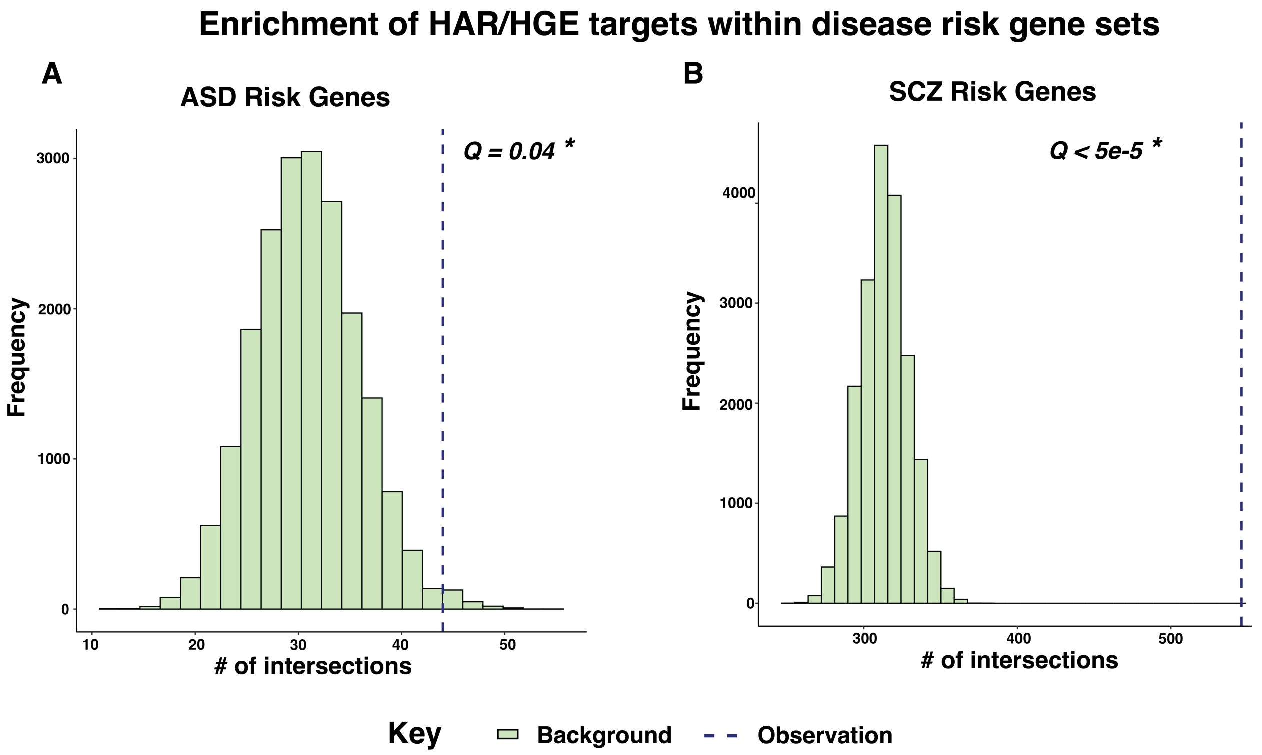


**Supplementary Figure 7. Enrichment of HAR and HGE gene targets within disease risk gene sets. (A)** Enrichment of HAR/HGE gene targets in the SFARI category I autism spectrum disorder (ASD) risk gene set [46]. *Q*-values were computed based on permutation using random sampling of the background (n = 20,000 trials; Methods). **(B)** Enrichment of HAR/HGE gene targets in the schizophrenia (SCZ) risk gene set [47]. *Q*-values were computed as in **(A).**

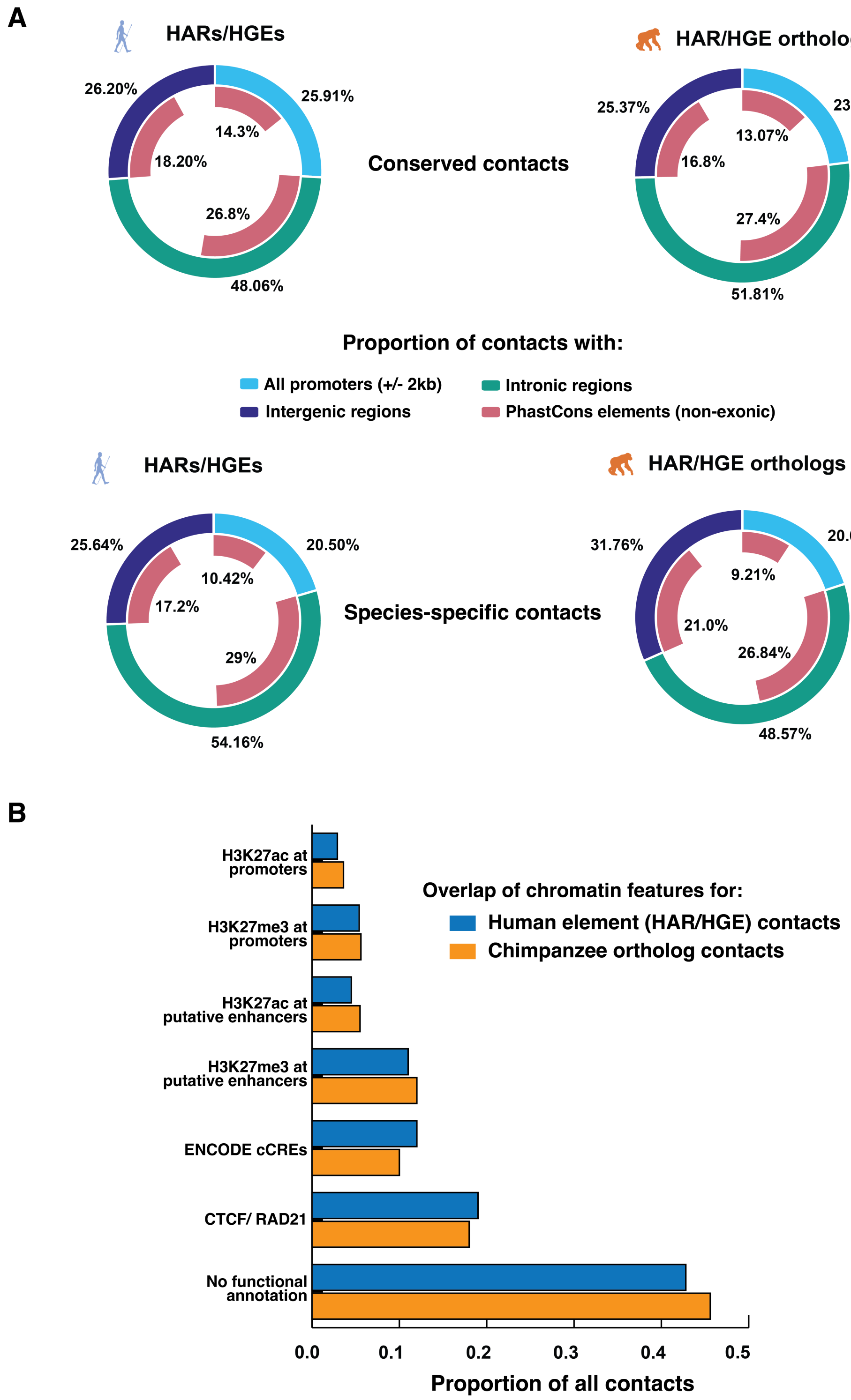


**Supplementary Figure 8. Functional properties of interactions involving HARs and HGEs.** **(A)** Doughnut plots showing the overlap of conserved and species-specific CHiC contacts with gene and phastCons conserved noncoding element annotations as described in the Results. **(B)** Bar plot showing the proportion of all CHiC interactions in hNSCs and cNSCs based on functional genomic data (H3K27ac, H3K27me3, CTCF, and RAD21 profiles generated for both species in this study and ENCODE cCREs).

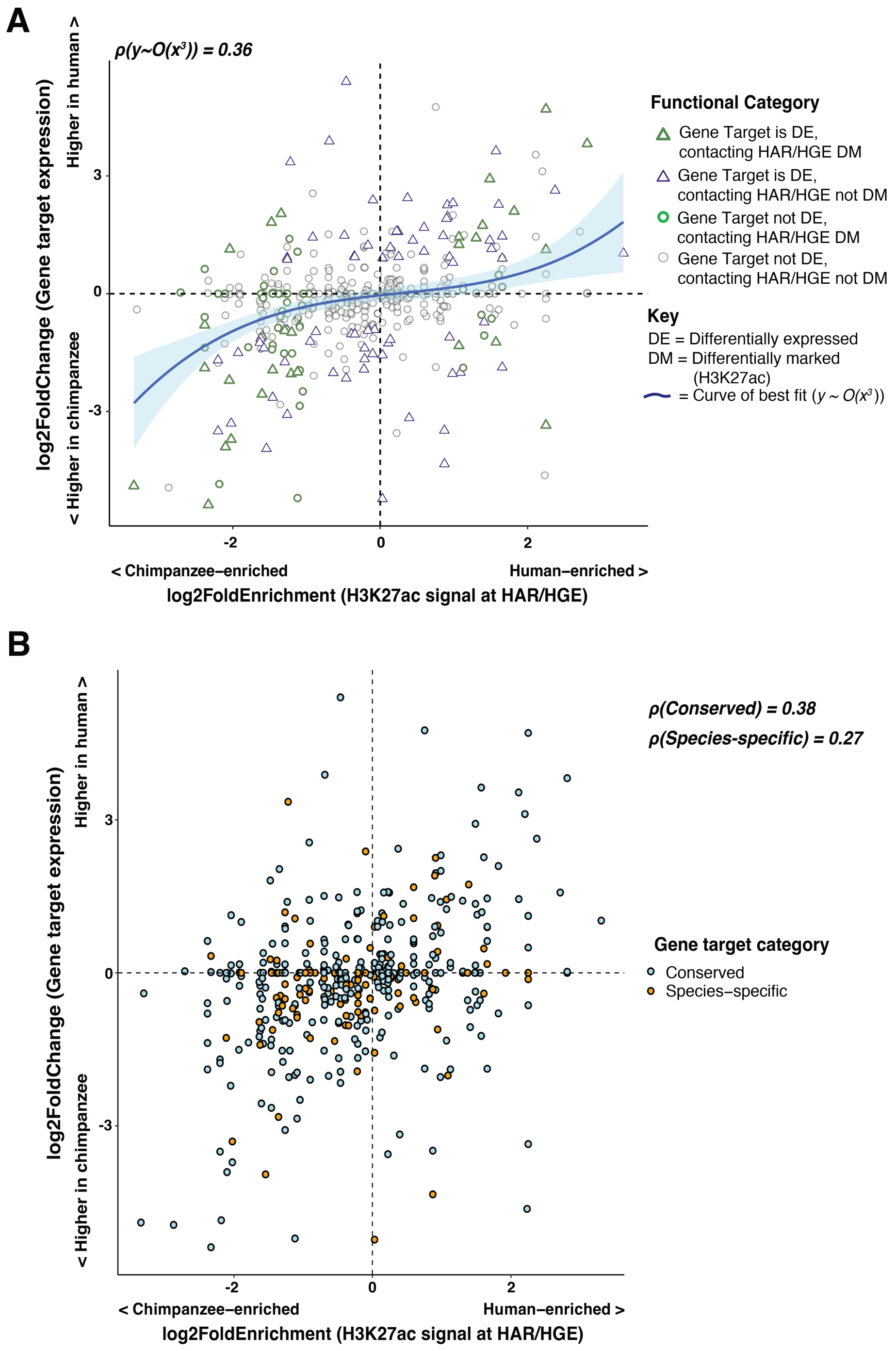


**Supplementary Figure 9. Association of relative expression for HAR and HGE gene targets against the relative H3K27ac level at the HAR or HGE (after Fig. 4A). (A)** Plot showing all combinations of HAR/HGE – gene target pairs: genes contacted by differentially marked (DM) elements are colored in green, and differentially expressed gene (DEG) targets are shown as triangles. The polynomial of best fit (y ~ O($x^3$)) is plotted in blue and standard error is shaded in light blue. **(B)** Plot where the combinations are colored by whether the gene target is conserved between hNSCs and cNSCs or is species-specific. Pearson correlation values (ρ) for each category are shown on the plot.

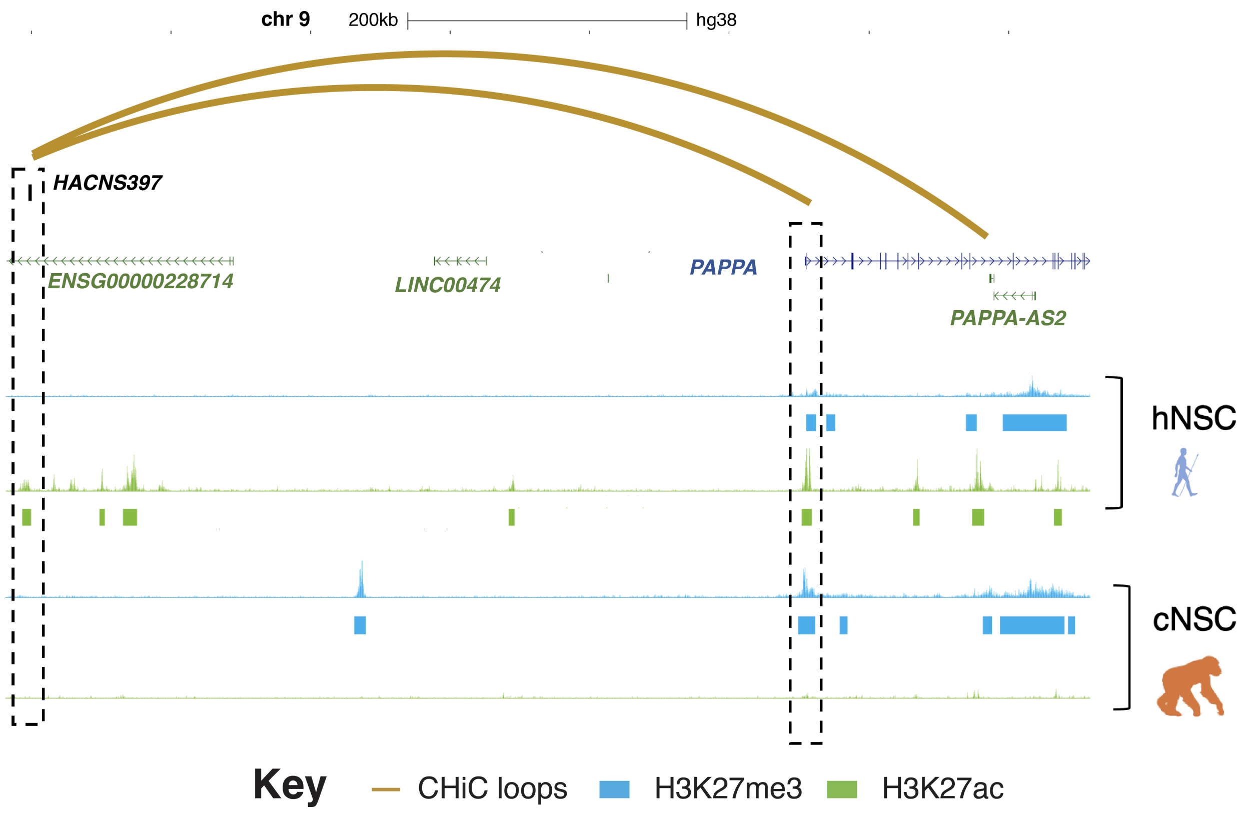


**Supplementary Figure 10. An example of a human-biased interaction.** *HACNS397* forms a loop differentially marked by H3K27ac in hNSCs compared to cNSCs (using DESeq2; Methods). Its gene target *PAPPA* is shown in blue, and curved golden lines indicate representative CHiC interactions involving the HAR. CUT&RUN signals are shown from top to bottom: H3K27me3 in blue and H3K27ac in green, for both hNSCs and cNSCs respectively. The overlap between CUT&RUN signal and the HAR and its gene target is highlighted by dashed black boxes.

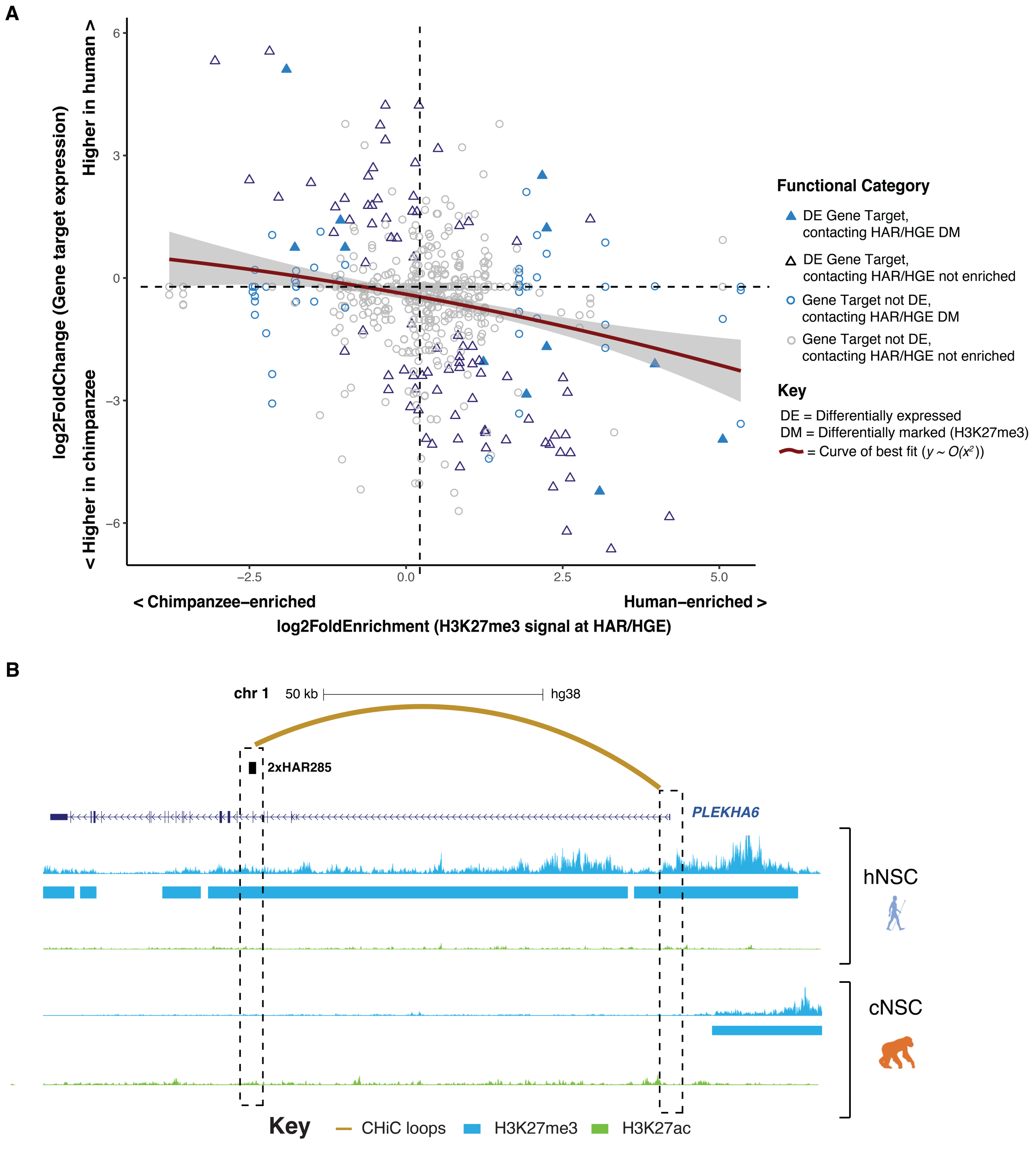


**Supplementary Figure 11. Differential H3K27me3 marking in hNSCs and cNSCs.** **(A)** Scatterplot of relative expression for all HAR and HGE gene targets (log2(human/chimpanzee)) against the relative H3K27me3 level at the HAR or HGE (log2(human/chimpanzee)). Genes contacted by differentially marked (DM) elements are colored in blue, and differentially expressed (DE) gene targets are shown as triangles. The polynomial of best fit (y ~ O($x^2$)) is plotted as a maroon curve and standard error is shaded in light grey. **(B)**

2xHAR285 forms a loop differentially marked by H3K27me3 in hNSCs compared to cNSCs (using DESeq2; Methods; Table S2). Its gene target *PLEKHA6* is shown in purple, and curved golden lines indicate representative CHiC interactions involving the HAR. The following CUT&RUN signal tracks are shown from top to bottom: H3K27me3 in blue and H3K27ac in green for both hNSCs and cNSCs, respectively. The overlap between CUT&RUN signal and the HAR and its gene target is highlighted by dashed black boxes.

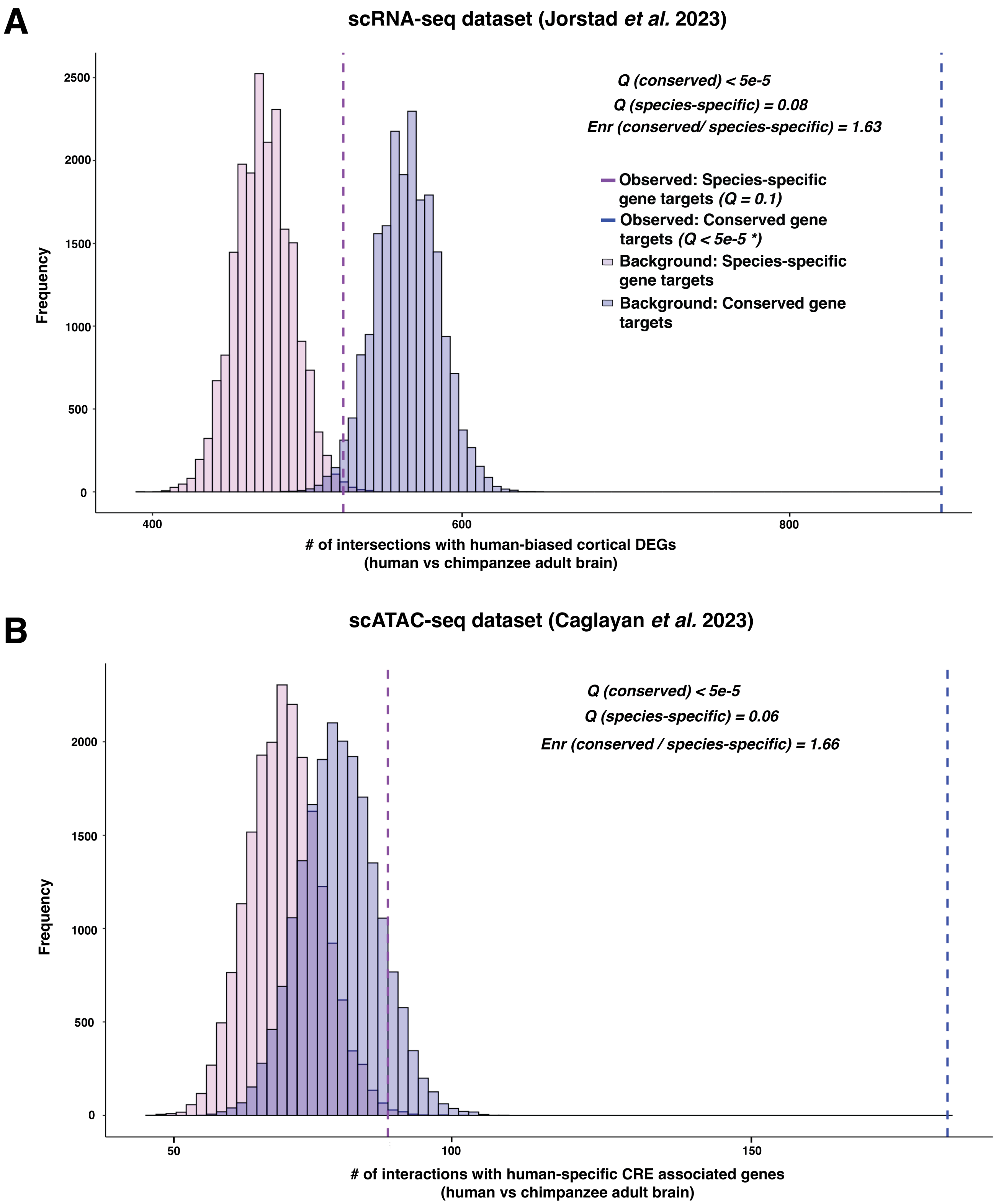
A
scRNA-seq dataset (Jorstad et al. 2023)
Q (conserved) < 5e-5
Q (species-specific) = 0.08
Enr (conserved/ species-specific) = 1.63
Observed: Species-specific gene targets (Q = 0.1)
Observed: Conserved gene targets (Q < 5e-5 *)
Background: Species-specific gene targets
Background: Conserved gene targets
Frequency
2500
2000
1500
1000
500
0
400
600
800
### of intersections with human-biased cortical DEGs
(human vs chimpanzee adult brain)
B
scATAC-seq dataset (Caglayan et al. 2023)
Q (conserved) < 5e-5
Q (species-specific) = 0.06
Enr (conserved / species-specific) = 1.66
Frequency
2000
1500
1000
500
0
50
100
150
### of interactions with human-specific CRE associated genes
(human vs chimpanzee adult brain)

**Supplementary Figure 12. Relative enrichment of conserved gene targets compared to species-specific gene targets among published gene sets identified through single cell sequencing. (A)** Enrichment of conserved HAR or HGE gene targets within human-biased DEGs called between human and chimpanzee adult cortex (scRNA-Seq data; Jorstad *et al* [34]), and their relative over-representation (abbreviated 'Enr') within the DEG set compared to species-specific gene targets. *P*-values were computed based on permutation using random sampling of the background followed by Bonferroni correction to get *Q* values (n = 20,000 trials; Methods). **(B)** Enrichment of conserved HAR or HGE gene targets in genes associated with cis-regulatory elements (CRE)- showing human-biased chromatin accessibility between human and chimpanzee cortex (scATAC-Seq data; Caglayan *et al* [35]), and their relative over-representation (abbreviated 'Enr') within the DE gene set compared to species-specific gene targets. *Q*-values were computed as in **(A)**.

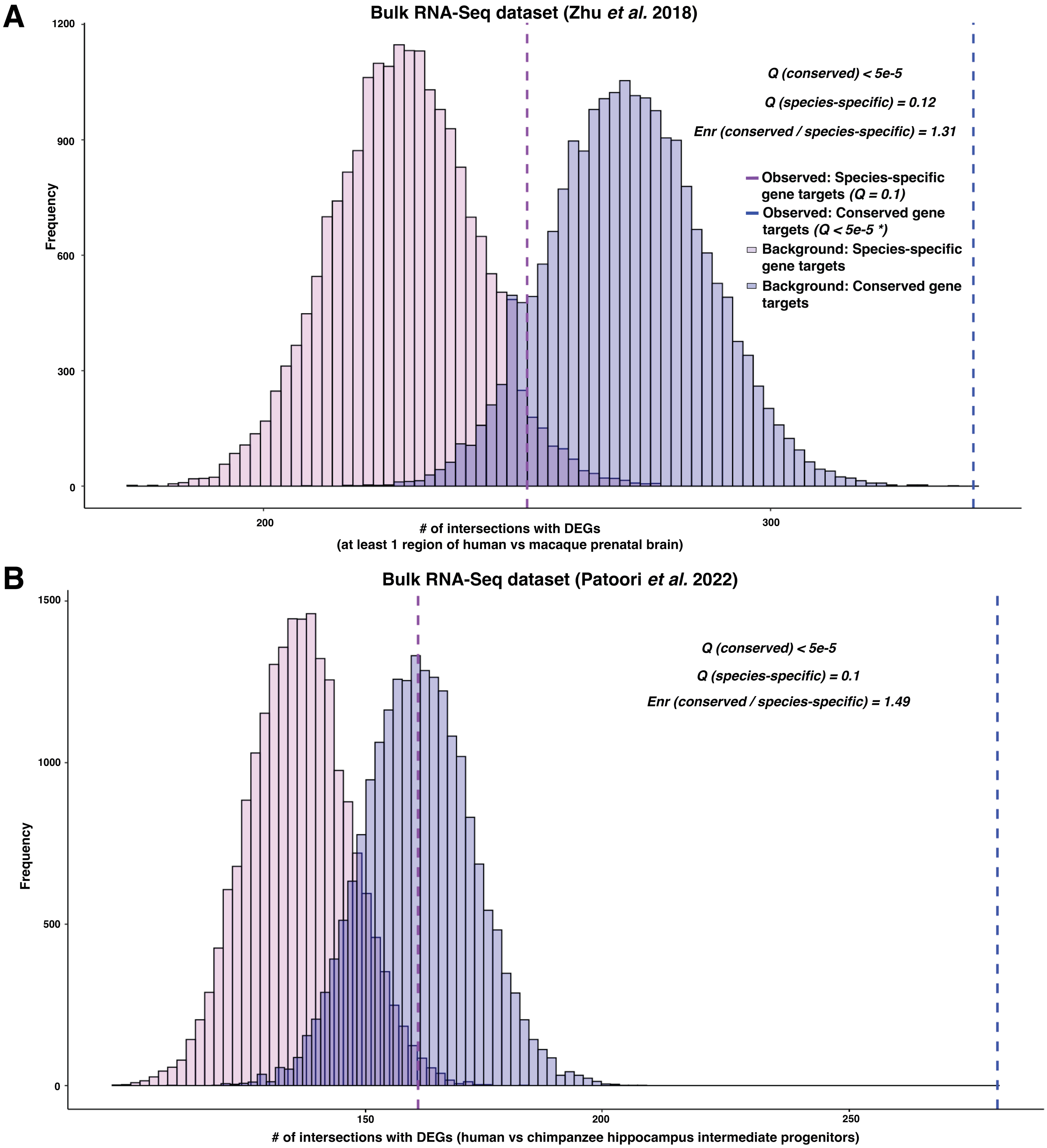
A
Bulk RNA-Seq dataset (Zhu et al. 2018)
Q (conserved) < 5e-5
Q (species-specific) = 0.12
Enr (conserved / species-specific) = 1.31
Observed: Species-specific gene targets (Q = 0.1)
Observed: Conserved gene targets (Q < 5e-5 *)
Background: Species-specific gene targets
Background: Conserved gene targets
Frequency
1200
900
600
300
0
200
300
### of intersections with DEGs
(at least 1 region of human vs macaque prenatal brain)
B
Bulk RNA-Seq dataset (Patoori et al. 2022)
Q (conserved) < 5e-5
Q (species-specific) = 0.1
Enr (conserved / species-specific) = 1.49
Frequency
1500
1000
500
0
150
200
250
### of intersections with DEGs (human vs chimpanzee hippocampus intermediate progenitors)

**Supplementary Figure 13. Relative enrichment of conserved gene targets compared to species-specific gene targets among published gene sets identified via bulk RNA-Seq. (A)** Enrichment of conserved HAR or HGE gene targets within DEGs called in at least one brain region for human versus macaque prenatal brain (bulk RNA-Seq data[36]), and their relative over-representation (abbreviated 'Enr') within the DE gene set compared to species-specific gene targets. *Q*-values were computed as in Fig. S12. **(B)** Enrichment of conserved HAR or HGE gene targets within DEGs called between cultured human and chimpanzee hippocampus intermediate progenitors[56], and their relative over-representation (abbreviated 'Enr') within the DE gene set compared to species-specific gene targets. *Q*-values were computed as in Fig. S12.

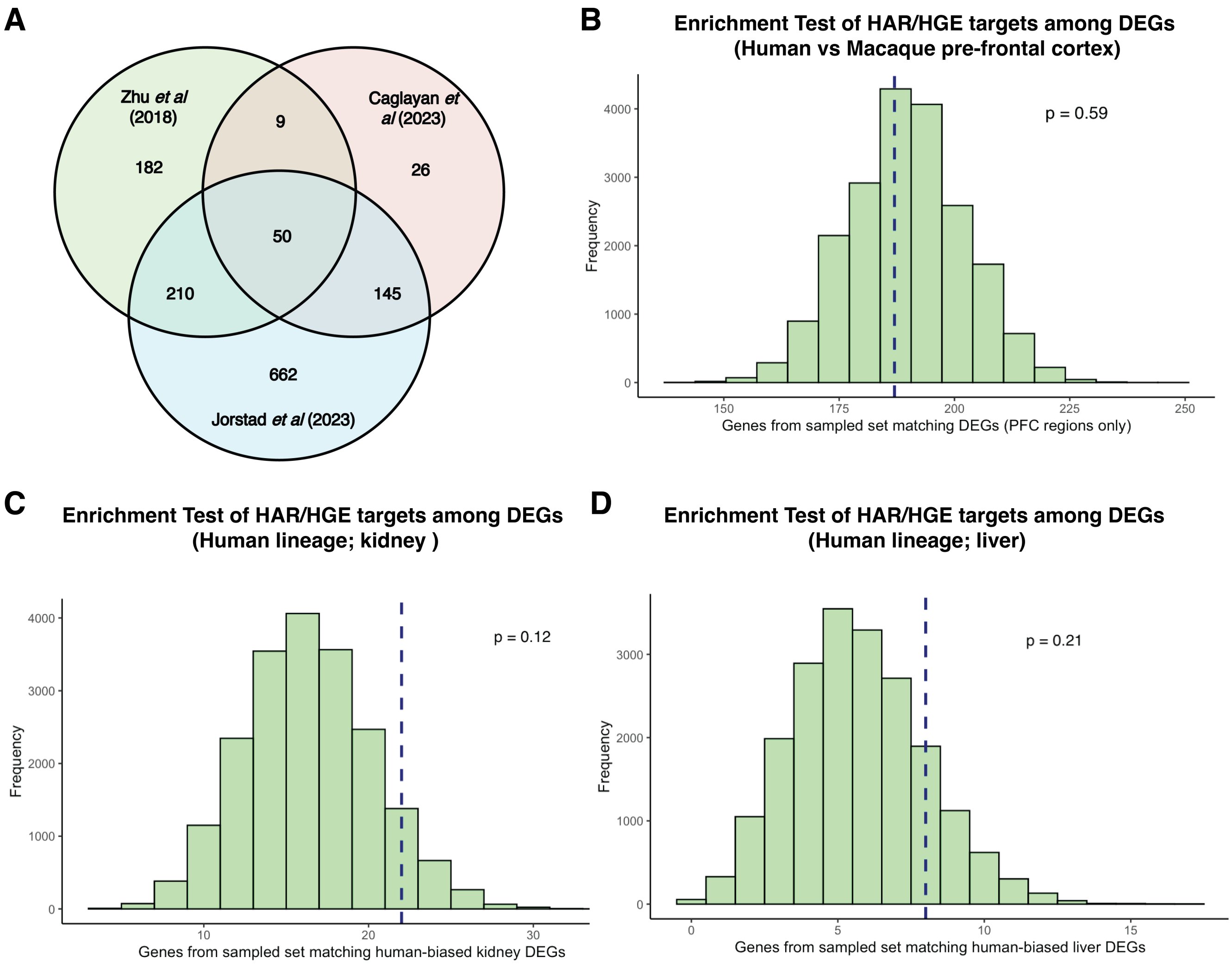


**Supplementary Figure 14. Enrichment of HAR and HGE gene targets among functional gene sets. (A)** Overlap of HAR/HGE gene targets in hNSCs which are called as human-specific DEGs from comparison of adult human and chimpanzee cortex (Jorstad *et. al* [34];blue bubble) with those called as DEGs between fetal human and Rhesus macaque brain (Zhu *et. al* [36]; green bubble) and genes associated with cis-regulatory elements (CRE)- showing human-biased chromatin accessibility between human and chimpanzee cortex (Caglayan *et. al* [35]; pink bubble). **(B)** HAR and HGE gene targets in hNSCs are not enriched within DEGs identified in a comparison of human and Rhesus macaque fetal pre-frontal cortex (data from Zhu *et. al* [36]) compared to other sampled protein-coding gene sets (Methods). *P*-values were computed based on permutation using random sampling of the background (n = 20,000 trials). **(C-D)** Enrichment test for HAR and HGE gene targets in gene sets which show lineage-specific expression shifts in the human lineage (compared to other primates/amniotes) compared to other sampled protein-coding gene sets, in kidney and liver respectively[57]. *P*-values were computed as in (**B**).

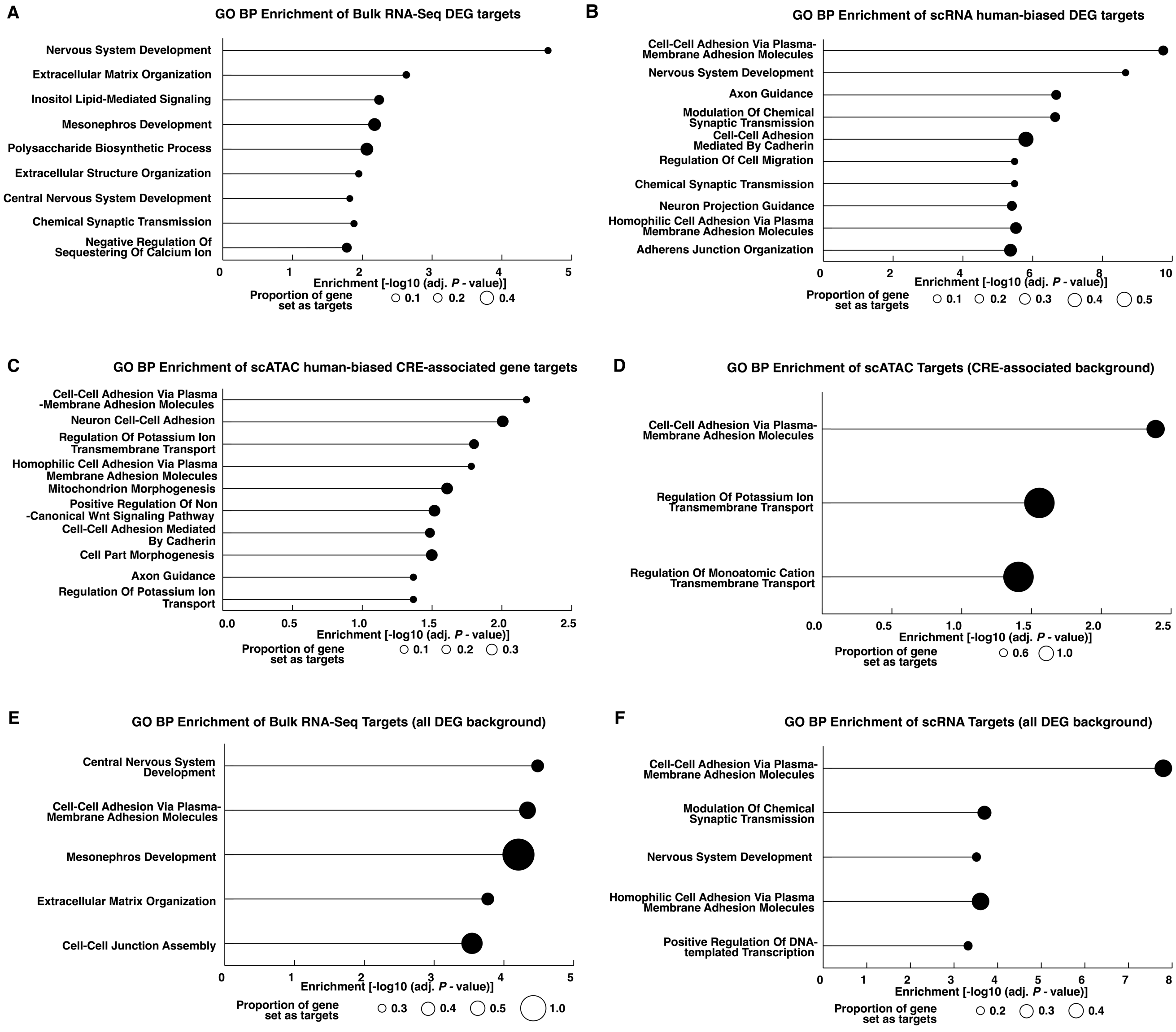


**Supplementary Figure 15. GO BP functional term enrichment for HAR and HGE gene targets among published gene sets. (A- C)** GO BP functional terms enriched for each of the HAR/HGE gene target subsets overlapping genes reported in each of the three published datasets in Fig. 4 (see also Fig. S14A). The background is all protein-coding genes. **(D- F)** GO BP functional terms enriched for HAR/HGE gene target subsets overlapping genes reported in each of the three published datasets in Fig. 4. The background is the full list of functional gene sets (DEGs or CRE-associated genes) from each of the studies.

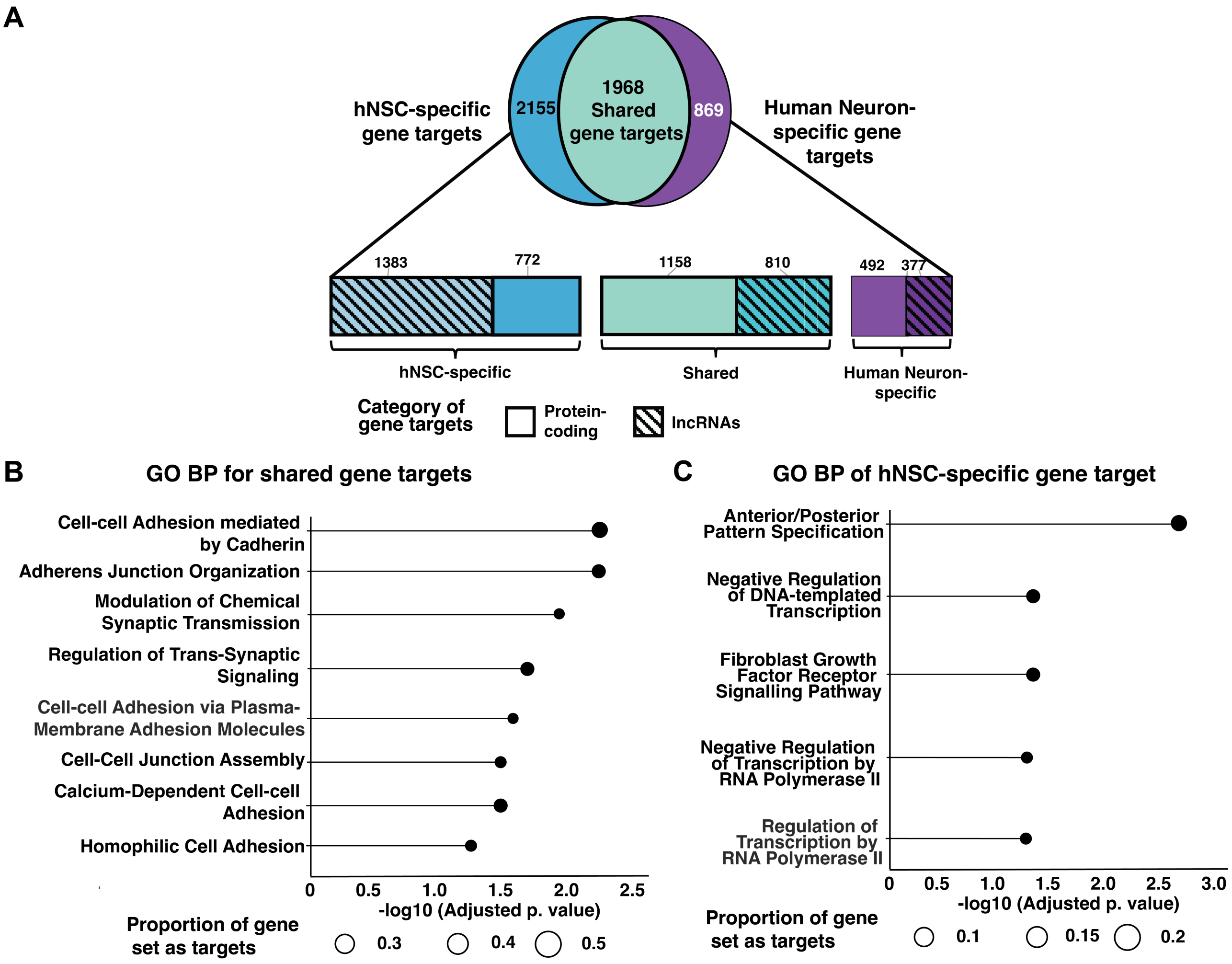


**Supplementary Figure 16**. **Gene targets of HARs and HGEs in neurons. (A)** Gene targets of HARs and HGEs in hNSCs vs neurons based on the human GENCODEv43 annotation as defined in the Results and Methods. The horizontal bar plot further subdivides hNSC-specific, neuron-specific, and shared targets into protein-coding and long non-coding RNA (lncRNA) genes. **(B)** GO Biological Process enrichment analysis performed on the shared set of protein-coding gene targets. **(C)** GO Biological Process enrichment analysis performed on the hNSC-specific set of protein-coding gene targets. The neuron-specific set is not shown, as there was no significant enrichment for any GO BP terms.

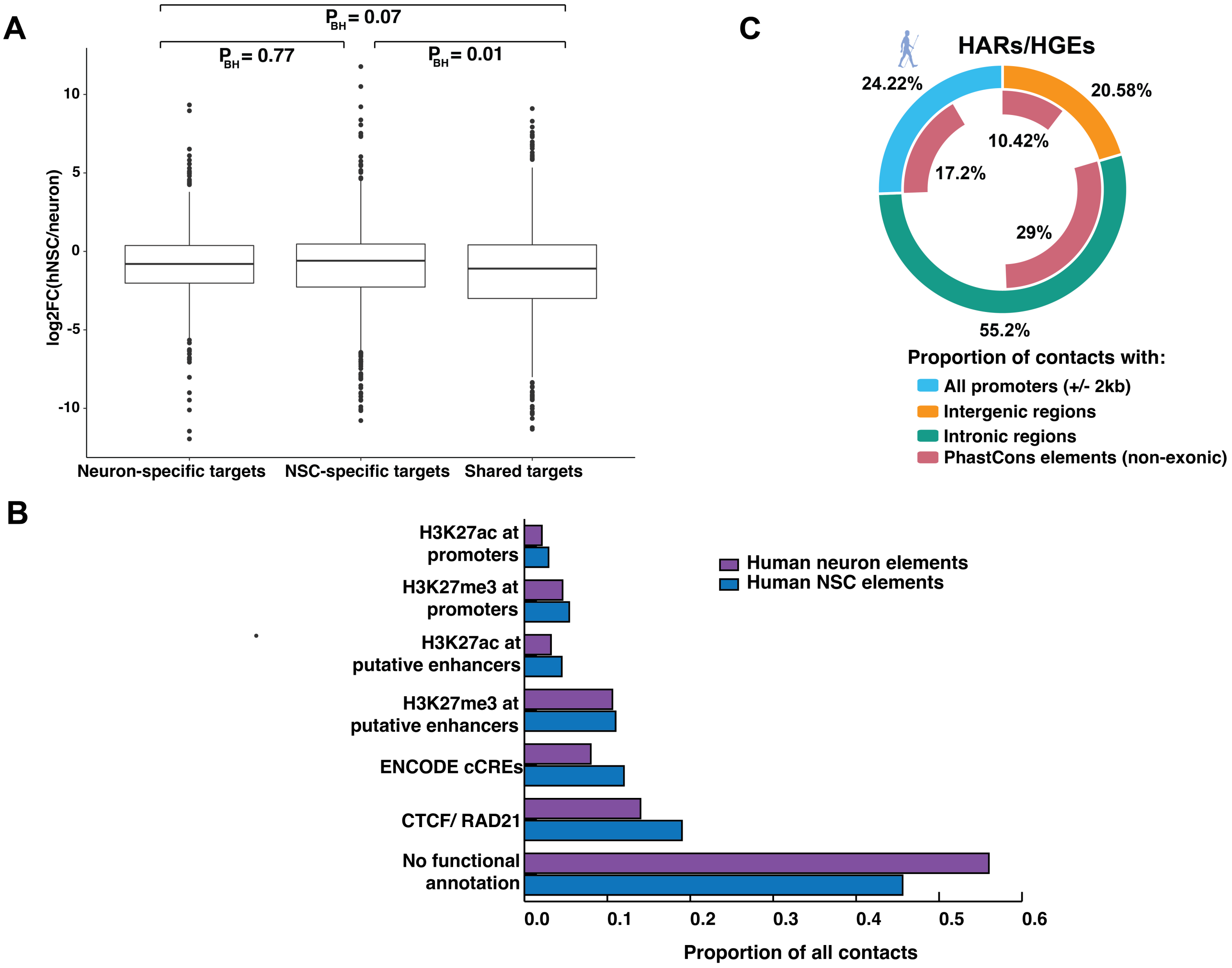


**Supplementary Figure 17**. **Functional annotation of interactions involving HARs and HGEs in neurons. (A)** Distribution of relative gene expression of gene targets shared between hNSCs and neurons versus cell type-specfic gene targets in hNSCs and human neurons (values provided are log2(hNSC TPM/neuron TPM). *P* values were computed from a Wilcoxon test and corrected using the BH procedure. **(B)** Doughnut plot showing the distribution of CHiC interactions in neurons based on gene and phastCons conserved noncoding element annotations as described in the Results. **(C)** Bar plot showing the proportion of CHiC interactions in neurons (purple) and hNSCs (blue) based on functional genomic data (H3K27ac, H3K27me3, CTCF, and RAD21 profiles generated in this study and ENCODE cCREs).

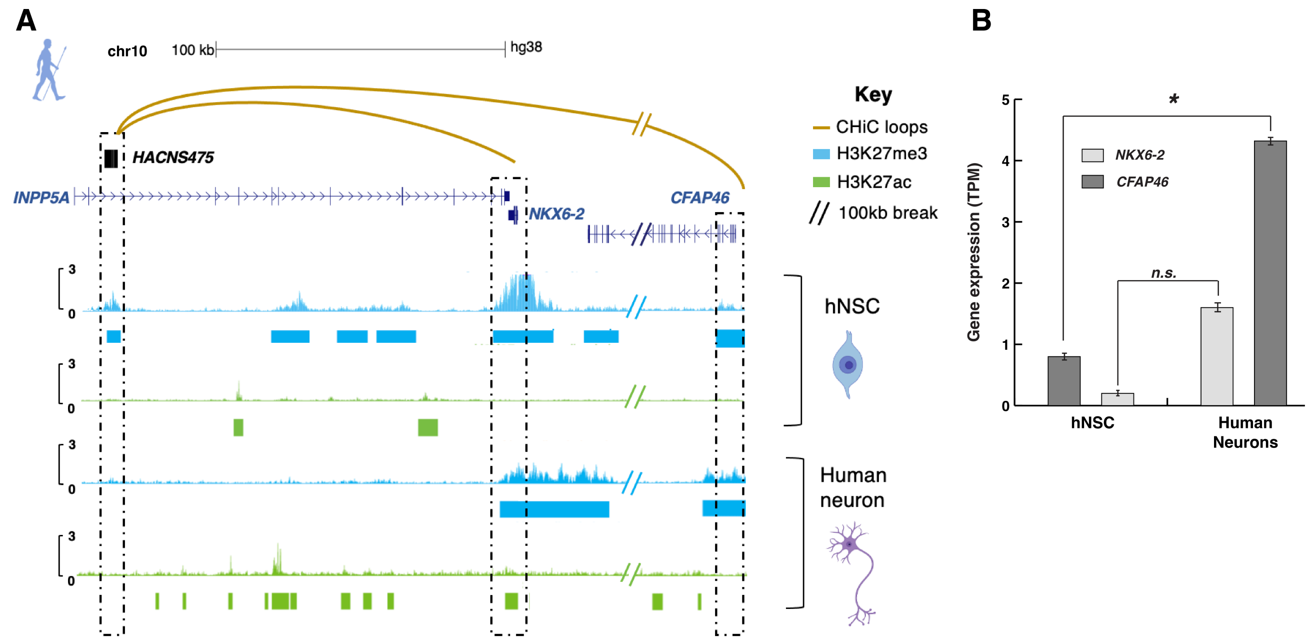


**Supplementary Figure 18**. **Example of a repressed loop in hNSCs that loses H3K27me3 marking upon differentiation. (A)** *HACNS475* forms a H3K27me3-marked loop with its target genes *NKX6-2* and *CFAP46* in hNSCs. In human neurons the H3K27me3 mark is present at the gene targets, but is lost at the HAR. Curved golden lines indicate significant CHiC interactions involving the HAR, and the following CUT&RUN signal tracks are shown from top to bottom: H3K27me3 in blue; H3K27ac in green for both hNSCs and hNSC-derived neurons. The overlap between CUT&RUN signal and HAR/target is highlighted by dashed black boxes. **(B)** Change in the expression of gene targets of *HACNS475*, *NKX6-2* and *CFAP46*, between hNSCs and neurons (TPM values, * = $P_{BH} < 0.05$; n.s. = not significant).

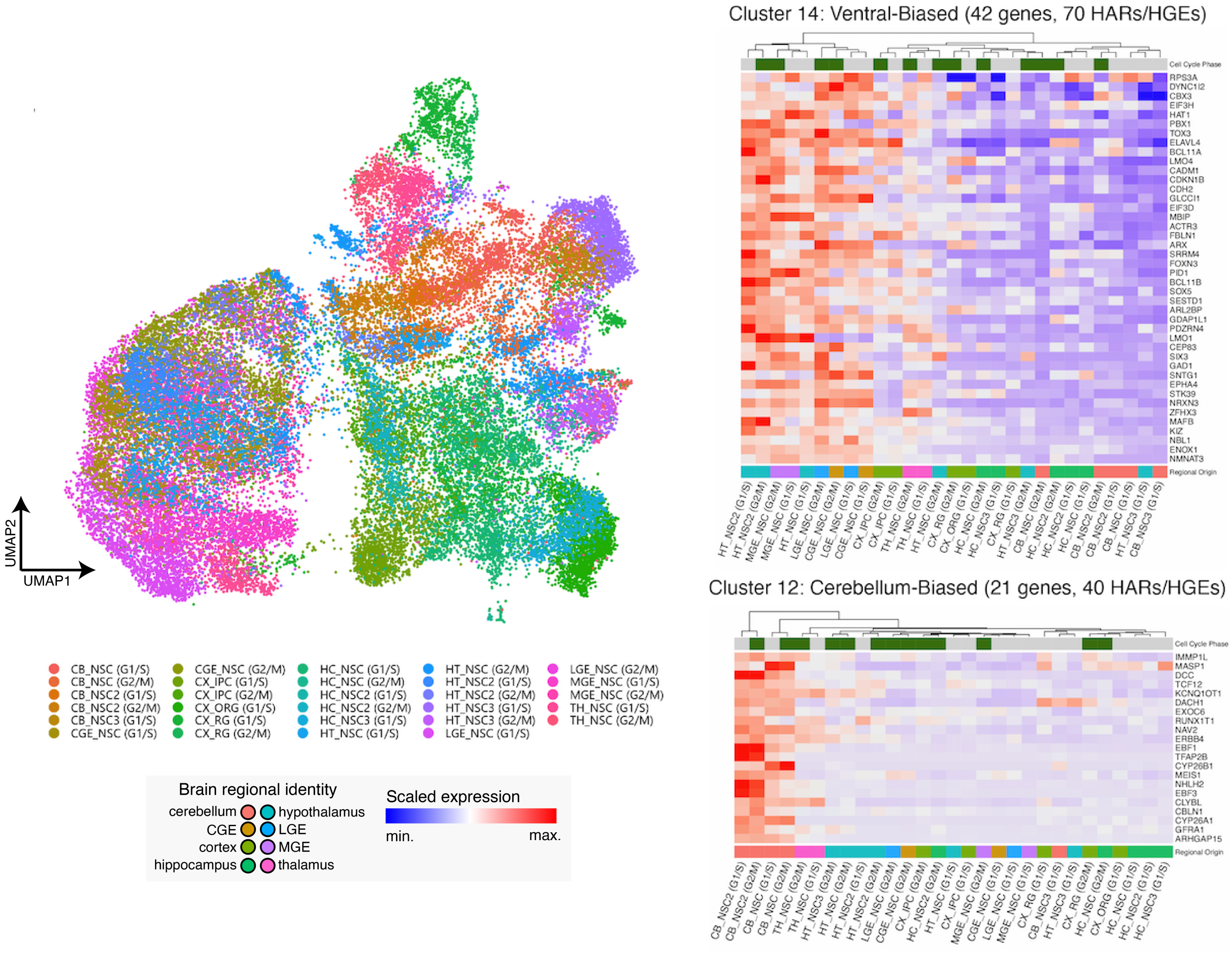


**Supplementary Figure 19. HAR and HGE gene targets identified in hNSCs show region-specific expression profiles in progenitors of the developing human brain.** *Left.* UMAP showing neural progenitors in scRNA-seq from eight embryonic and fetal human brain regions colored by cell type. *Right*. Heatmaps showing clustered average expression profiles of HAR/HGE gene targets across neural stem cell populations in the cerebellum and ventral telencephalon/diencephalon. Brain regions are labeled as in Fig. 6.

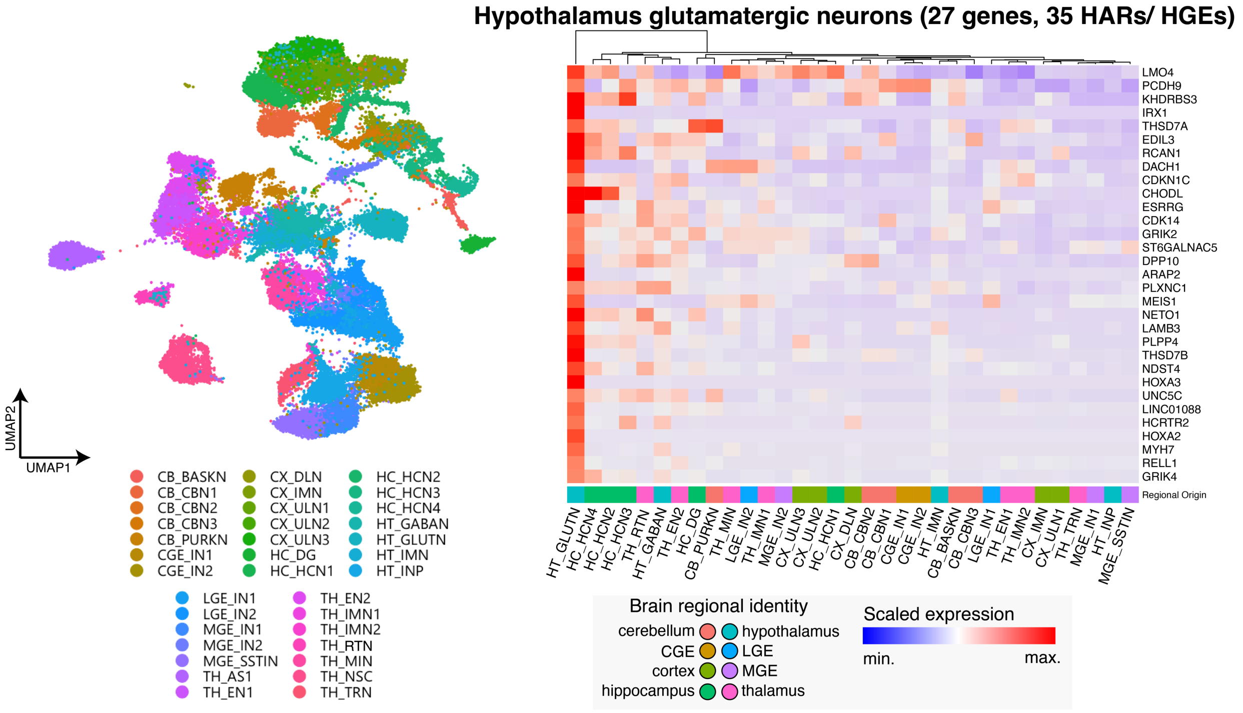


**Supplementary Figure 20. HAR and HGE gene targets identified in human neurons show region-specific expression profiles in neurons of the developing human brain.** *Left.* UMAP showing neurons in scRNA-seq from eight embryonic and fetal human brain regions colored by cell type. *Right*. Heatmap showing the clustered average expression profiles of HAR/HGE gene targets across specific neuron subtypes; in this case, hypothalamus glutamatergic neurons, represented by the UMAP on the left as HT_GLUTN. Other abbreviations for cell types called in neurons of the fetal brain are explained in Table S3.

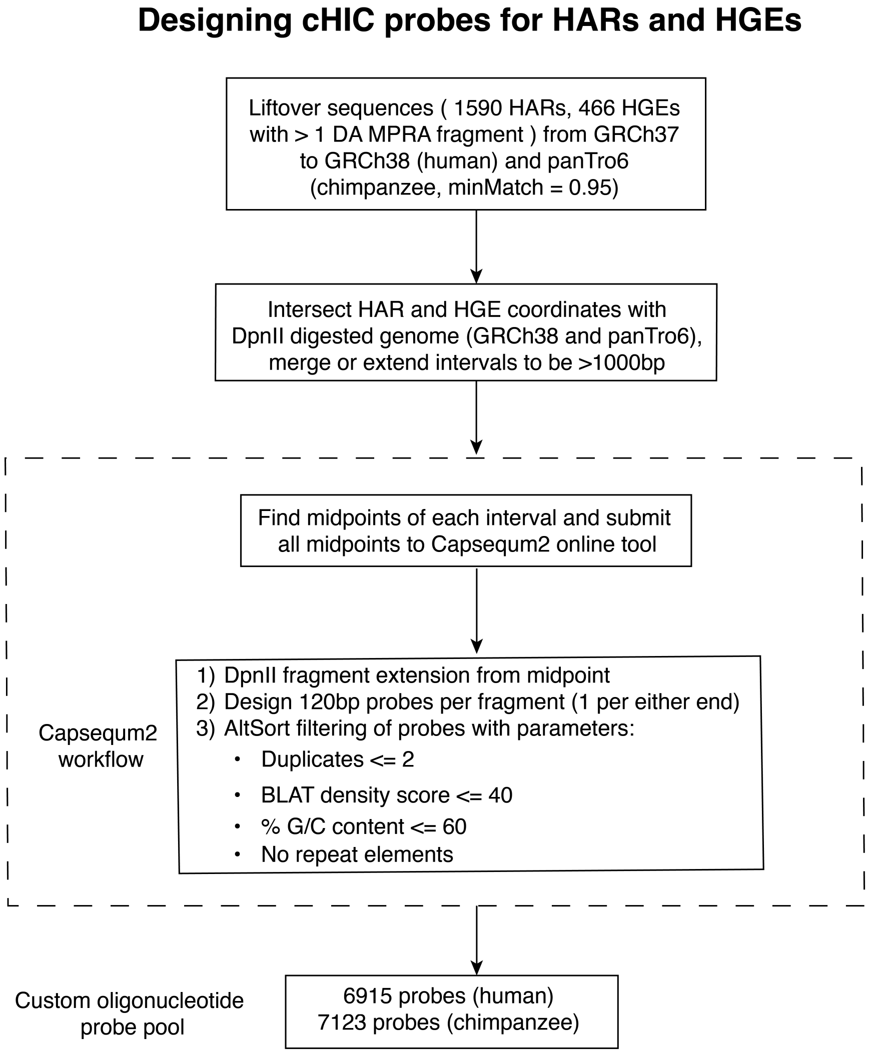


**Supplementary Figure 21. Probe design for CHiC study.** Flow chart describing the criteria and filters used in designing biotinylated probes for the capture step of HAR and HGE interactions in the human and chimpanzee CHiC assays.

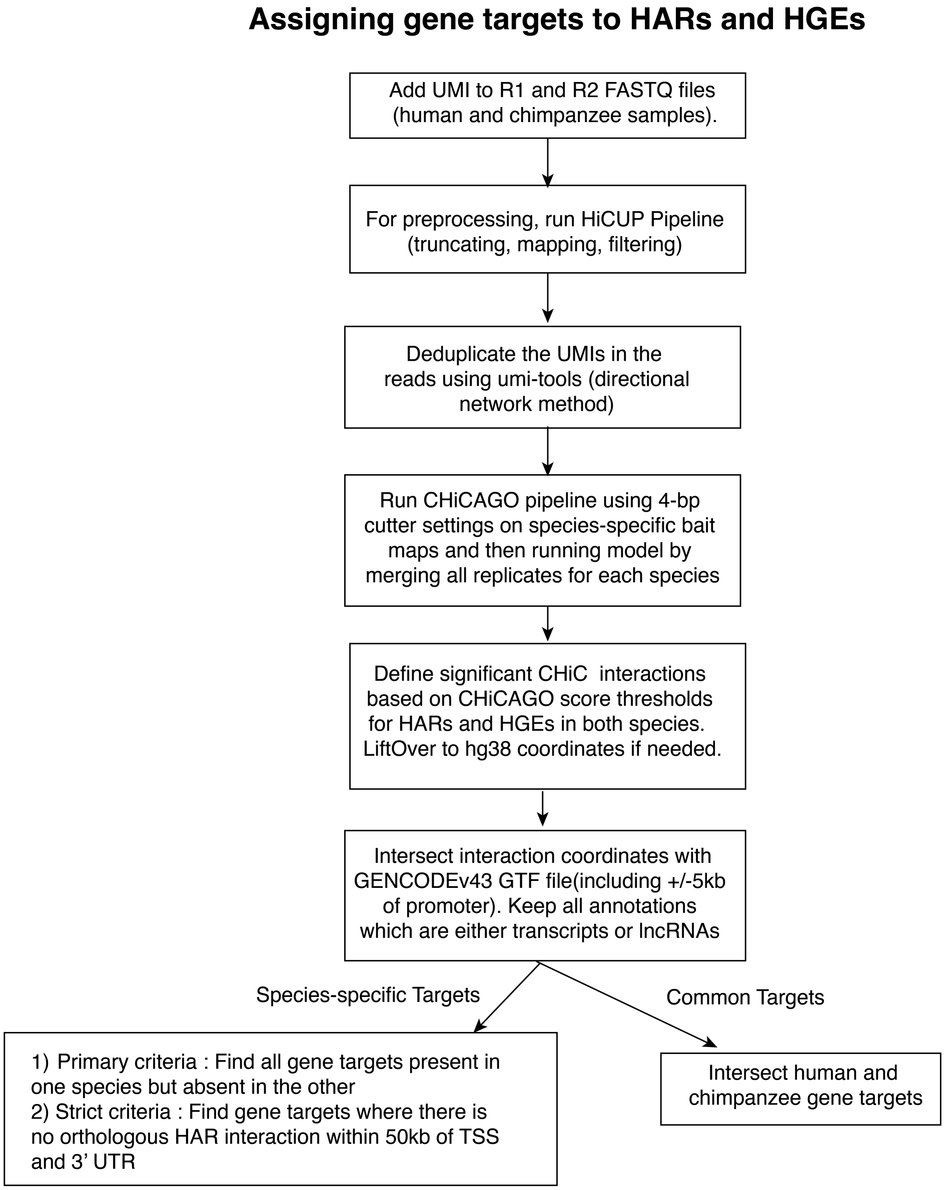


**Supplementary Figure 22. Identification of gene targets of HARs and HGEs.** Flow chart describing the criteria used to link significant CHiC interactions to gene targets (and logic behind splitting them into conserved and species-specific gene target categories).
